# Pericentrin is a Kinesin-1 Activator that Drives Centriole Motility

**DOI:** 10.1101/2022.01.12.476023

**Authors:** Matthew R. Hannaford, Rong Liu, Neil Billington, Zachary T. Swider, Brian J. Galletta, Carey J. Fagerstrom, Christian Combs, James R. Sellers, Nasser M. Rusan

## Abstract

Centrosome positioning is essential for their function. Typically, centrosomes are transported to various cellular locations through the interaction of centrosome nucleated microtubules with motor proteins. However, it remains unknown how centrioles migrate in cellular contexts in which centrioles do not nucleate microtubules. Here, we demonstrate that during interphase inactive centrioles move directly along the non-centrosomal microtubule network as cargo for the motor protein Kinesin-1. We identify Pericentrin-Like-Protein (PLP) as a novel Kinesin-1 interacting molecule essential for centriole motility. PLP directly interacts with the cargo binding domain of Kinesin-1 and they comigrate on microtubules *in vitro*. Finally, we demonstrate that PLP-Kinesin-1 dependent transport is essential for centrosome asymmetry age-dependent centrosome inheritance in asymmetric stem cell division.

## Introduction

Centrosomes are organelles comprised of two centrioles surrounded by a matrix of proteins termed the pericentriolar material (PCM). The PCM recruits gamma-tubulin ring complexes to centrosomes in a cell cycle dependent manner to create a microtubule organizing center (MTOC; Azimzadeh and Bornens, 2007). Centrosomes are important for organizing cilia and mitotic spindles, both of which rely on proper centrosome positioning (Tang and Marshall, 2012).

Most research on centrosome movement has focused on the separation of centrosomes during late G2 prior to mitosis (Tanenbaum and Medema, 2010; Agircan et al., 2014). Separation is coordinated by motor proteins exerting pushing and pulling forces on centrosomal microtubules (MTs). The key motor protein involved in this process is Kinesin-5/Eg5, which acts on antiparallel MTs to slide them in opposing directions (Kapitein et al., 2005), and dynein located at the cell cortex and the nuclear envelope which exerts pulling forces on the MTs (Dujardin and Vallee, 2002). Together these motors position centrosomes for bipolar spindle formation. Incomplete centrosome separation prior to NEB can result in the formation of merotelic kinetochore-MT attachments (Silkworth et al., 2012). Merotelic attachments lead to lagging chromosomes and chromosome instability. In severe cases, failure to separate centrosomes can result in supernumerary centrosomes in the following cell cycle. This can lead to spindle abnormalities that further promote chromosome instability(Ganem et al., 2009); a common feature among cancer cells (Bakhoum and Cantley, 2018).

Importantly, centrioles are also motile during interphase, and this motility is poorly understood. In multi-ciliated cells hundreds of centrioles are produced which must then migrate to the cell cortex to function as basal bodies (Spassky and Meunier, 2017; Jord et al., 2019; Ching et al., 2021). If centriole migration is impaired, then cilia formation is effected, which can result in a range of disorders termed ciliopathies (Reiter and Leroux, 2017). Interestingly, despite the importance of basal body positioning at the cell cortex, little is known about how they reach their destination (Dawe et al., 2006). Centrioles have also been observed to be highly motile prior to cytokinesis, with a proposed function at the midbody to regulate abscission (Piel et al., 2001; Krishnan et al., 2021).

One of the most documented instances of interphase centriole motility occurs in *Drosophila* neuroblasts (NBs). In NBs the centrioles are highly asymmetric in PCM levels. At mitotic exit, the daughter centriole recruits the protein Centrobin which is simultaneously shed by the mother centriole (Januschke et al., 2011, 2013; Gallaud et al., 2020). The localization of Centrobin to the daughter centriole precedes PCM recruitment and MTOC formation. It is through this MT-nucleation capacity that the daughter centriole is stably localized to the apical side of the interphase NB resulting in a polarized MT network (Rebollo et al., 2007; Rusan and Peifer, 2007; Januschke and Gonzalez, 2010). Meanwhile, the mother centriole sheds PCM (Conduit and Raff, 2010; Ramdas Nair et al., 2016), resulting in a centriole incapable of MTOC activity, it then moves from the apical to the basal side of the cell throughout interphase. As the NB enters the next prophase, this mother centriole once again recruits PCM in preparation for bipolar spindle formation (Rusan and Peifer, 2007; Rebollo et al., 2007). Analysis of mutant NBs revealed that the centriolar proteins Pericentrin-like-protein (PLP; Lerit and Rusan, 2013) and Bld10/Cep135 (Singh et al., 2014), the motor protein Kinesin-1, and the Kinesin-1 activator Ensconsin/Map7 (Gallaud et al., 2014; Métivier et al., 2019) are involved in centriole separation. However, the underlying mechanism regulating centriole movement remains unclear.

In this study we take advantage of *Drosophila* to examine the mechanism of centriole motility. We show that centrioles are transported along MTs by Kinesin-1, via a direct interaction of the Kinesin Heavy Chain (KHC) with PLP. We propose that Kinesin-1 and PLP work together to ensure mother centriole motility in NBs. The consequence of failed Kinesin-PLP driven motility is defective centrosome asymmetry and errors in age-dependent centriole segregation.

## Results

### Inactive centrioles are highly motile in interphase

During interphase, *Drosophila* centrosomes shed their PCM, leaving behind a centriole which does not nucleate microtubules (MTs), we will refer to these centrioles as inactive. Inactive centriole dynamics are best characterized in the *Drosophila* neuroblasts (NBs) where the mother centriole is inactivated in the late stages of mitosis and upon entry into interphase, migrates from the apical side of the cell towards the basal side (Figure 1A,D; Movie 1). The high temporal resolution imaging required for accurate quantitative analysis of centriole movement is challenging in NBs due to the rapid speed of centrioles and three-dimensional nature of motility. Therefore, we sought to identify non-NB cell populations with inactive, motile centrioles that would be suitable for our study. Inactive centrioles are found in other cell populations(Rogers et al., 2008) and we found that both *Drosophila* S2 cells (Figure 1B,E; Movie 2) and the squamous epithelial cells that constitute the peripodial membrane of the imaginal discs (McClure and Schubiger, 2005), hereon referred to as peripodial cells (PCs; Figure 1C,F; Movie 3) contained inactive and highly motile centrioles. The flat nature of the PC layer allowed for tracking multiple centrioles simultaneously with high temporal resolution. We confirmed that the motile centrioles in NBs, S2 cells, and PCs did not recruit the PCM component Centrosomin (Cnn), indicating they are indeed inactive during interphase (Figure 1D-F).

**Figure 1:**
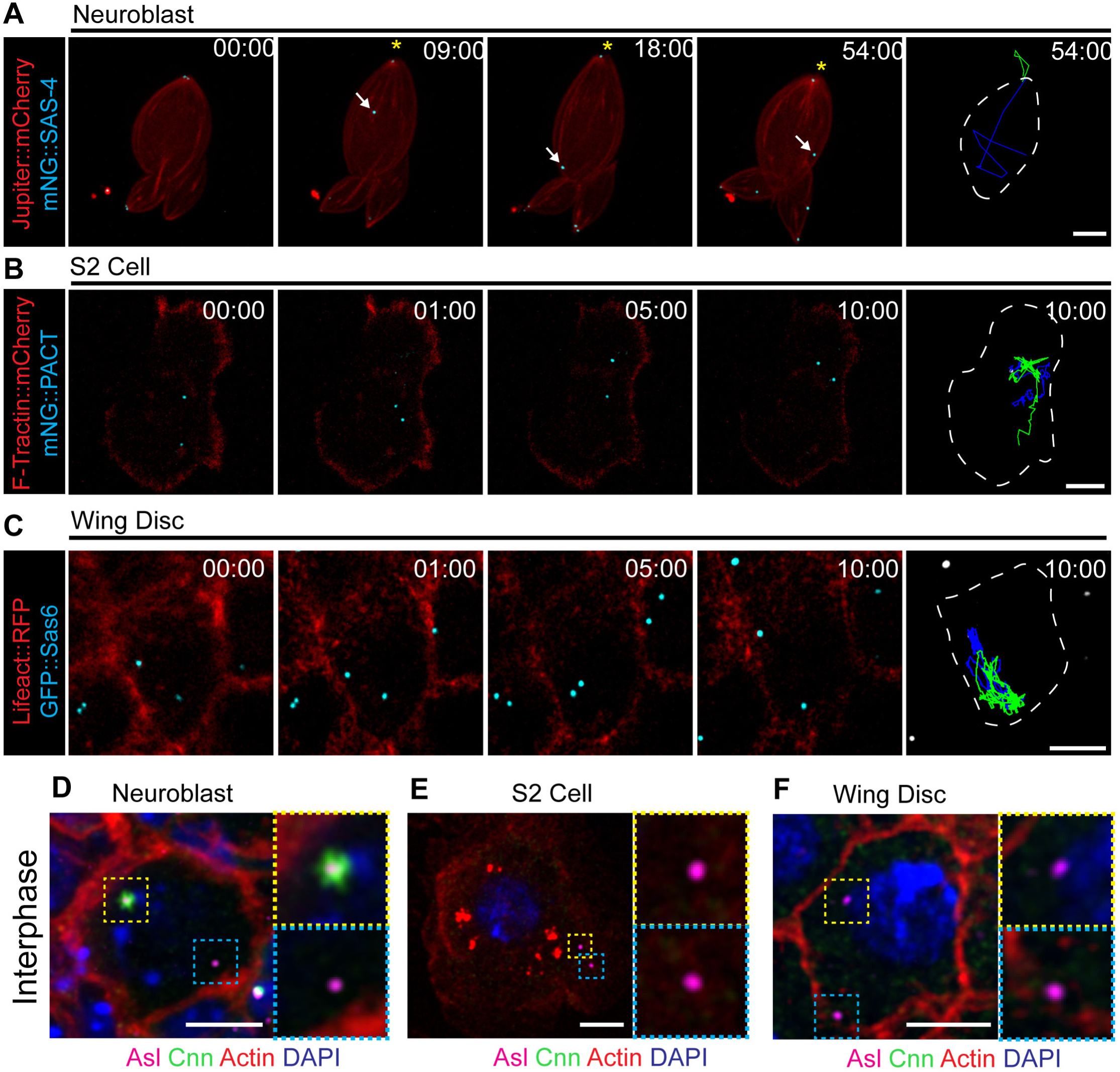
*Drosophila c*entrioles are motile in interphase cells. **A)** Z-stack projection of an interphase NB expressing Jupiter::mCherry (red) and mNG::Sas4 (cyan). The mother centriole (asterisk) remains closely associated with the apical cell cortex; the daughter centriole (arrow) moves throughout the cell. **B)** Cultured S2 cell transfected with F-tractin::mCherry (red) to visualize the cell and mNG::PACT (cyan) to label centrioles. Both centrioles are highly motile through the acquisition. **C)** Peripodial cell expressing Lifeact::RFP (red) to visualize the cell and GFP::Sas6 (cyan) labelling the centrioles. Both centrioles are highly motile within the cell. Last column in A, B and C show cell outline (white line) and a 10 minute time projections of centriole movement (green and blue lines). **D)** Fixed NB showing that Cnn (Green) is restricted to one of the two centrioles (magenta). **E)** Fixed S2 cell and **F)** Peripodial Cell showing no Cnn (Green) present on the centrioles (magenta). Scale bars: 5µm. Time Stamp: mm:ss.

### Inactive centrioles are MT cargo

The precise mechanism of centriole motility is unknown, but both the MT and actin cytoskeleton have been implicated in centriole movement and positioning (Piel et al., 2000; Burakov et al., 2003). Using Latrunculin-A to depolymerize the F-Actin network in PCs, we found that an intact actin network was not required for centriole movement (Figure S1; Movie 4). To investigate the role of the MT cytoskeleton, we depleted MTs by incubating wing discs on ice and then allowing them to recover in Colcemid containing media (Figure 2A; Movie 5). Tracking centrioles in MT depleted PCs revealed a near complete loss of motility with average instantaneous velocity and mean square displacement approaching zero (Figure 2B,F,G; Movie 5). We conclude that the MT network is essential for centriole motility. To test if dynamic MTs were required for motile centrioles, we treated wing discs with Colchicine in the absence of ice, which did not disassemble MTs (Figure 2C), but did block MT polymerization (growth) as revealed by the lack of EB-1 localization (Figure 2D). This treatment resulted in a 30% decrease in average instantaneous velocity but did not ablate centriole movement (Figure 2E-G; Movie 5). Therefore, MTs are necessary for centriole movement, but their nucleation and dynamics are not.

**Figure 2:**
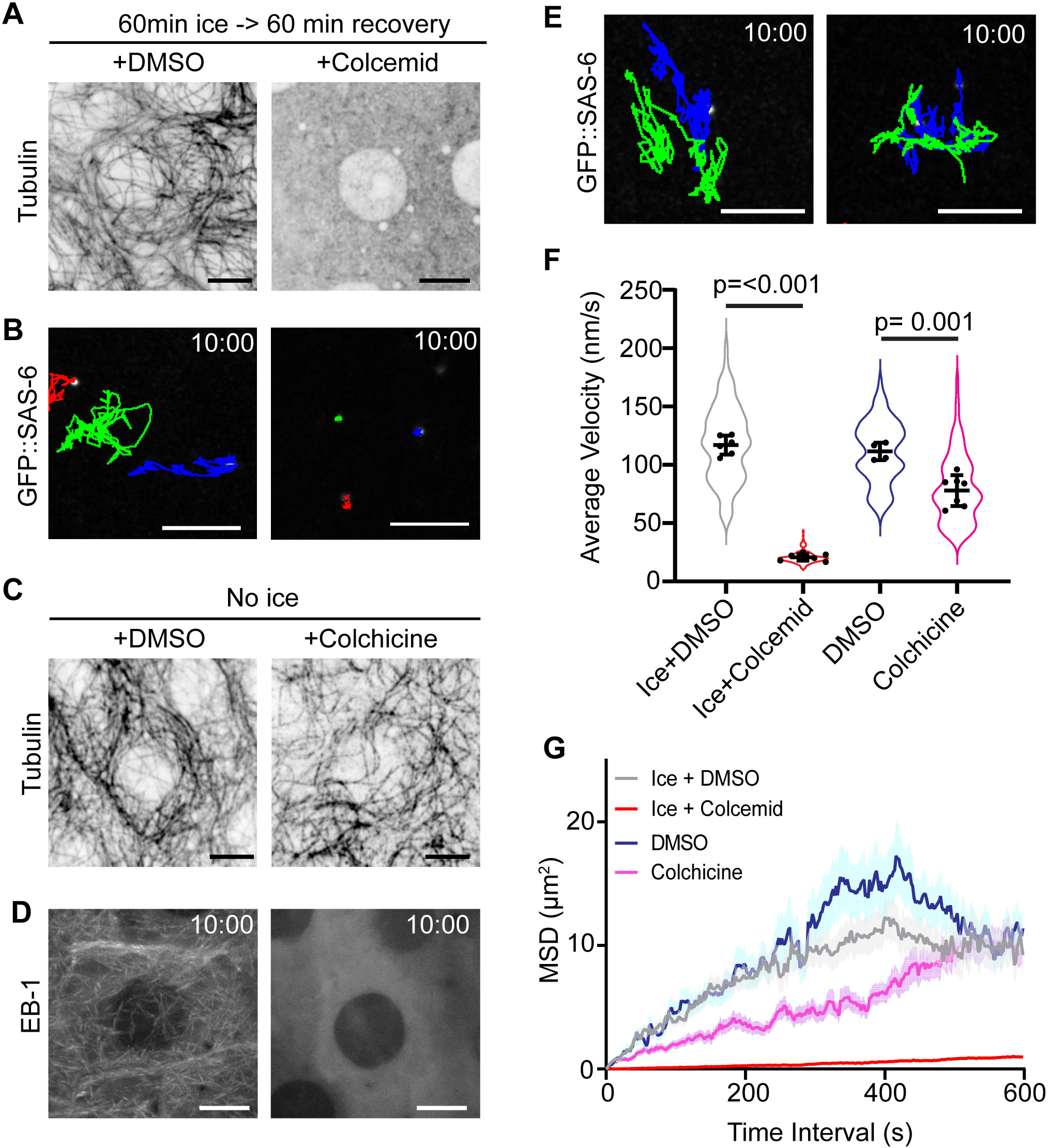
Interphase motility is dependent on intact microtubule networks. **A)** Peripodial cells following 1 hour ice treatment followed by recovery in DMSO or 50µM Colcemid. Note no visible microtubules remaining in the Colcemid treated wing disc. **B)** 10 minute time projection of centriole movement (colored tracks). Centrioles are not motile following ice treatment and Colcemid recovery. **C and D)** Colchicine treatment does not destroy the pre-existing microtubule network (C) but does block microtubule dynamics revealed by EB-1 localization (D). **E)** 10 minute time projections of centriole movement (colored tracks). Centrioles remain highly motile following Colchicine treatment. **F)** Quantification of instantaneous velocity in the indicated conditions (Ice + DMSO: 117±8.1 n= 6 wing discs, 95 centrioles. Ice+Colcemid: 20.7± 2.8, n=7 wing discs, 158 centrioles. DMSO: 115.5±7.4, n=4 wing discs, 78 centrioles. Colchicine: 77.9±13.2, n=7 wing discs, 109 centrioles). Data = Mean ± Standard Deviation, p values derived from unpaired t. test. **G)** Average mean squared displacement of centrioles. Scale bars: 5µm. Time stamp: mm:ss.

To further explore the relationship between centrioles and the MT network, we performed super resolution microscopy. Fixed imaging of MTs and centrioles in PCs and NBs revealed a close positional relationship between centrioles and MTs (Figure 3A,B), while live imaging revealed centrioles moving along MTs (Figure 3C,D; Movie 6,7). To robustly and easily label both centrioles and MTs in live PCs, we used the MT dye SIR-Tubulin, which revealed centrioles moving on MTs, even changing directions by switching their MT tracks (Figure 3E, S2; Movie 8). These data show that centrioles likely travel on MTs as cargo.

**Figure 3:**
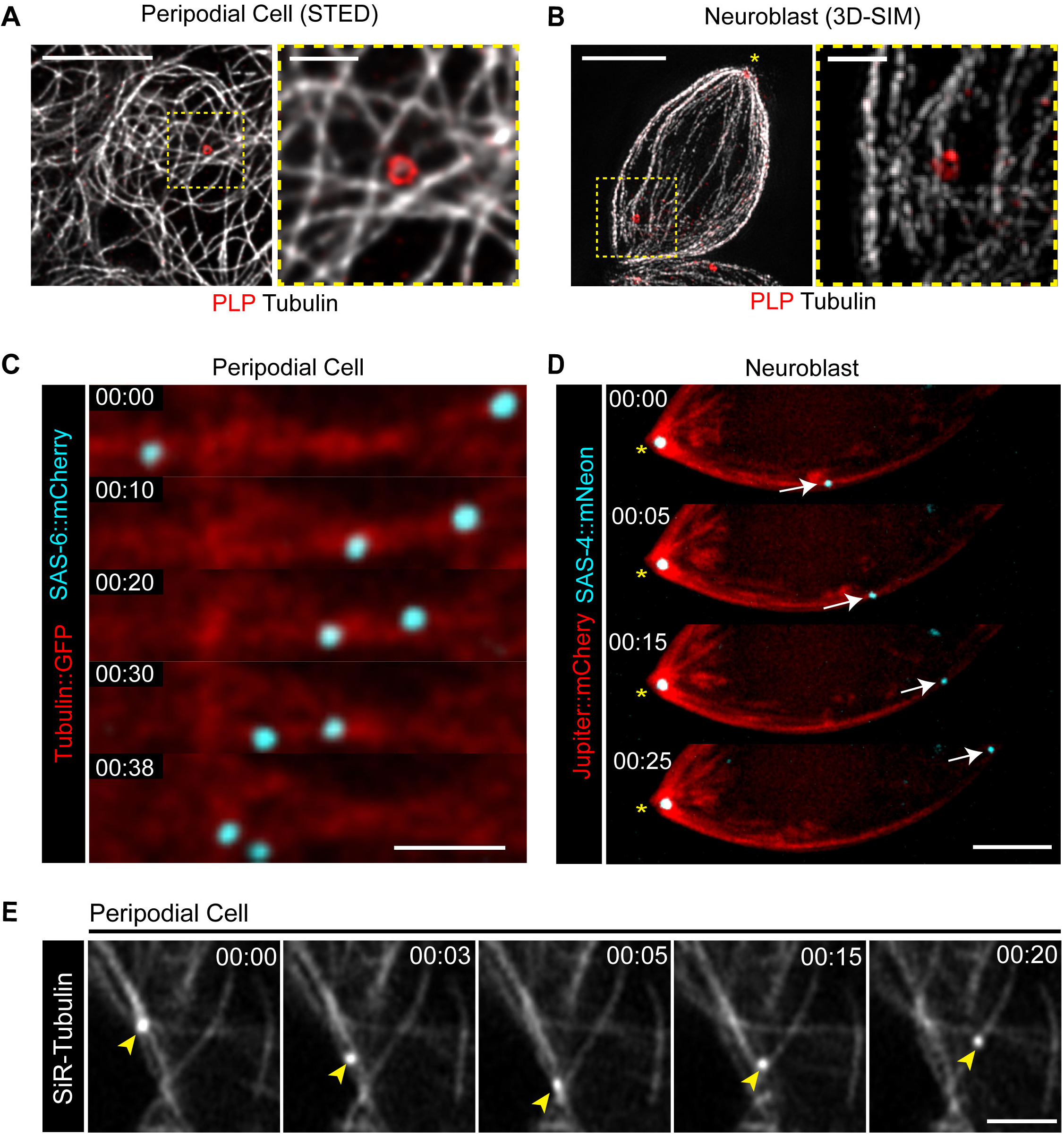
Centrioles are microtubule cargo. **A and B)** Super-resolution imaging reveals close relationship between centrioles (PLP; red) and microtubules (Gray) in PCs (A) and NBs (B). Scale bars: 5µm, inset 1µm. **C and D)** Centrioles (Cyan) move on the microtubules (red) in PCs, scale bar: 2µm (C) and NBs (D). Fluorescence transgenes are as indicated on left. **E)** PCs treated with SiR-Tubulin showing centrioles (arrowheads) moving along microtubules and switching tracks. Scale bar 2µm. Time stamp: mm:ss.

### Kinesin-1 is required for efficient centriole transport

To investigate the MT transport of centrioles, we aimed to identify which motor proteins were required for their motility. Previously, the heavy chain of *Drosophila* kinesin-1 (Khc) and its activator Microtubule Associated Protein 7/Ensconsin (Map7, Ens) were shown to be important for centrosome separation at prophase during asymmetric NB division (Gallaud et al., 2014; Métivier et al., 2019). Therefore, we tested if Kinesin-1 was involved in MT mediated centriole transport by knocking down Khc, Kinesin light chain (Klc), or Ens in PCs using the PC specific driver AGIR-GAL4 (Gibson et al., 2002). Control centrioles moved rapidly around the PCs as expected while knockdown of Khc, Klc or Ens significantly reduced centriole motility (Figure 4A-C; Movie 9). Thus, Kinesin-1 is important for interphase centriole motility.

**Figure 4:**
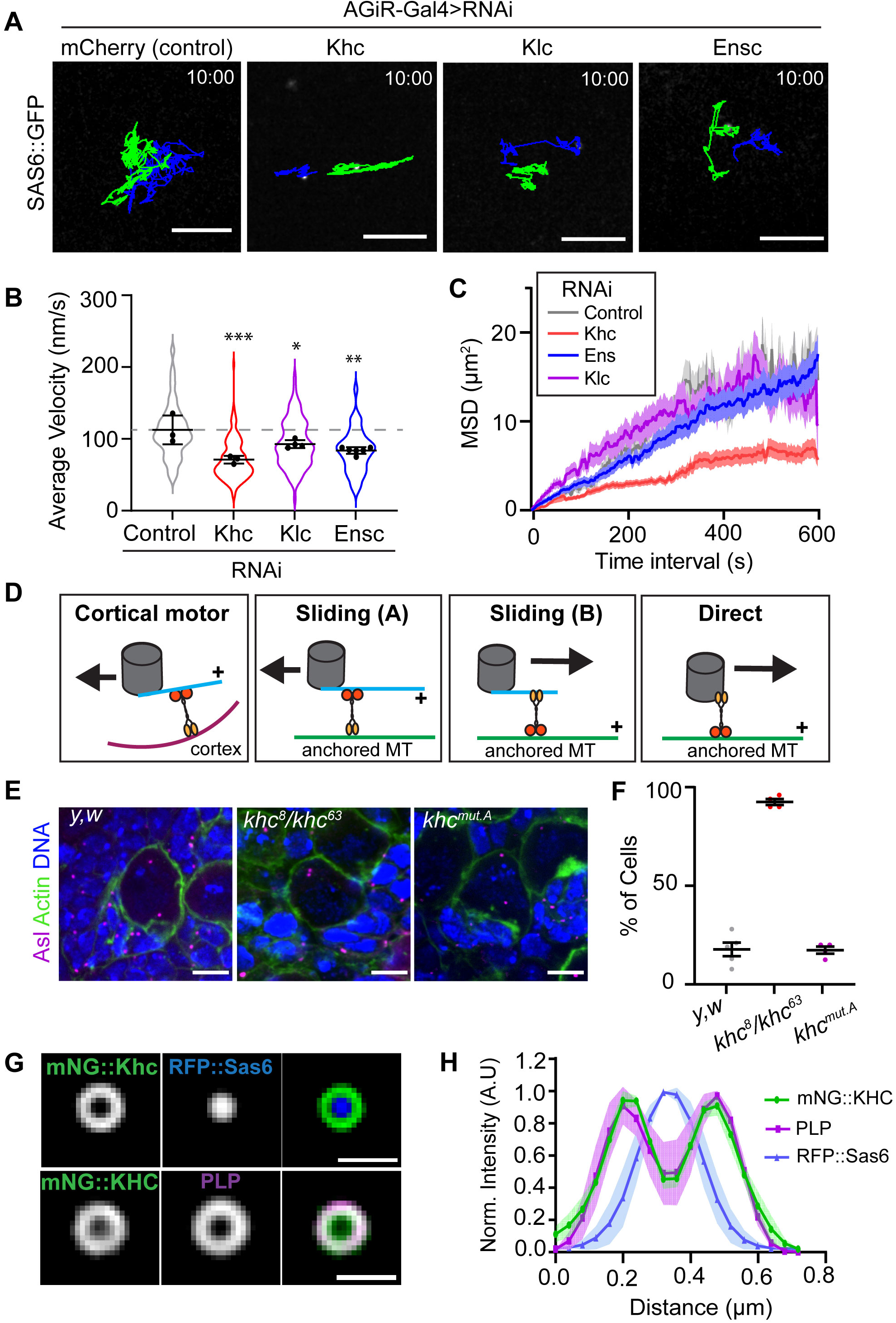
Kinesin-1 is required for efficient centriole motility. **A)** 10 minute time projections of centriole movement (colored tracks) in the indicated knockdown conditions in PCs. Scale bars: 5µm. **B)** Average velocity is significantly reduced following the knockdown of Kinesin-1 components (Control: 112±20, n=3 wing discs, 100 centrioles. KHCRNAi: 71±5, n=3 wing discs, 110 centrioles. KLCRNAi: 92± 5, n=4 wing discs, 97 centrioles. ENSRNAi: 83±8, n=6 wing discs, 84 centrioles; ANOVA p = 0.001, Dunnett’s pairwise comparisons: Ctrl vs. KHCRNAi p= 0.0005, ***, Ctrl vs KLCRNAi p = 0.043, *, Ctrl vs EnsRNAi p=0.0027, **). **C)** Mean squared displacement is reduced following KHC knockdown in PCs. **D)** Diagram summarizing the models by which Kinesin-1 could move centrioles in cells. Kinesin-1 cargo domain is shown in yellow; motor domain (orange) always walks toward the indicated + sign and the black arrow indicates movement direction of centriole. **E)** Z-stack projections of fixed NBs sowing centriole positioning. Note in *khc^8^/khc^63^* NBs the centrioles are adjacent at the apical side of the cell. **F)** Quantification of the percentage of neuroblasts with adjacent apical centrioles. y,w: 17.8% ± 7, n=5 brains; *khc^8^/khc^63^*: 92.5% ± 3, n=4 brains, *khc^mut.A^*: 17.4% ± 3.5, n=4 brains. **G)** Averaged OMX-SIM micrograph showing mNG::KHC localizes to the outer centriole edge. Scale bar: 500nm. **H)** Quantification of rotational averaged centrioles showing the distribution of mNG::KHC relative to PLP and RFP::Sas6 (n=4).

Based on these data, we hypothesized that Kinesin could move centrioles via three possible mechanisms. First, by indirect motor transport whereby the centriole nucleates a small number of MTs, that in-turn contact anchored Kinesin motor at the cortex (Figure 4D). We do not favor this model as, unlike Dynein, Kinesin-1 is not known to act in this manner. In addition, this model requires dynamic MTs that search the cortex for motors (such as cortical Dynein); however, our Colchicine treatment showed that MT nucleation and dynamics are not required for inactive centriole motility (Figure 2E). We also note that previous studies on inactive centrioles did not identify the presence of centriolar MTs in interphase *Drosophila* cells(Rusan and Peifer, 2007; Rebollo et al., 2007; Rogers et al., 2008).

The second possible mechanism of centriole motility is via the MT “sliding” activity of Kinesin-1 (Winding et al., 2016; Lu et al., 2016) (Figure 4D, Sliding A and B). KHC slides MTs by anchoring its C-terminal to one MT via a binding site within the cargo binding domain and walking along a second MT with its motor domain. This sliding activity can be reduced by 50% using a *khc* allele in which C-terminal MT binding site is mutated (*khc^mut.A^*: R914A, K915A, R916A, and Q918A(Lu et al., 2016; Winding et al., 2016)). To test whether sliding was essential for centriole motility we compared centriole positioning between control, *khc^8^/khc^63^* (hypomorphic), and *khc^mut.A^* NBs (Figure 4E). As expected, the *khc^8^/khc^63^* NBs exhibited a failure to separate their centrioles (Gallaud et al., 2014). However, reducing the sliding activity of Kinesin-1 had no effect on centriole separation compared to controls (Figure 4F). Additionally, either sliding model (Figure 4D, Sliding A and B) would require that both the centriole attached MTs and the anchored MT to be precisely polarized relative to each other and relative to the apical-basal cell axis in order to generate the required directional motility in NB. We believe this would be extremely difficult to achieve, and thus do not favor a sliding model.

Finally, the third possible mechanism involves Kinesin-1 directly binding the centriole surface and moving it as cargo. For this model to be plausible, Kinesin-1 would have to localize to centrioles. Kinesin-1 has previously been shown to localize to centrosomes in vertebrate cultured cells (Neighbors et al., 1988) but its localization to centrosomes in *Drosophila* has not been investigated. To examine Kinesin-1 localization we expressed and imaged mNeon::KHC in *Drosophila* S2 cells. Structured illumination microscopy revealed that mNeon::KHC localized as a ring around RFP::Sas-6 (core centriole marker), while occupying the same spatial position as Pericentrin-like-protein (PLP; Figure 4G,H). These data indicate that KHC is positioned on the outer layer of the centriole in a region occupied by proteins that bridge the centriole and PCM in mitosis. Collectively, our data thus far suggest a model whereby Kinesin-1 moves centrioles as cargo by directly binding the centriole surface.

### The centriole bridge protein PLP is required for centriole motility

PLP is a recruiter and organizer of PCM, and to perform this function it localizes to the outer edge of the centriole (Varadarajan and Rusan, 2018). Importantly, previous mutant analysis has implicated PLP in centrosome positioning in NBs as well as basal body positioning in sensory neurons (Martinez-Campos et al., 2004; Lerit and Rusan, 2013; Galletta et al., 2014; Roque et al., 2018). However, the mechanism by which PLP modulates PCM levels and influences centriole/centrosome motility remains unknown. Based on the localization of PLP to the outer layer of the centriole and PLP’s role in centriole/centrosome positioning, we hypothesized that PLP is required for MT-based motility of centrioles in interphase cells.

To test our hypothesis, we knocked down PLP in PCs using two independent RNAi lines. In each case, PLP knockdown caused a substantial 70% reduction in centriole motility (Figure 5A-C; Movie 10). The loss of centriole motility could be explained in two ways. Firstly, it is possible that PLP depletion resulted in ectopic PCM recruitment, similar to what was previously described in NBs (Lerit and Rusan, 2013), leading to the formation of centrosomal MTs that anchor the centrosome to the cortex. However, the lack of PCM on centrioles in PLP RNAi PCs excludes this possibility (Figure S3). Therefore, we favor a second explanation that PLP is directly involved in centriole motility, potentially functioning as a Motor-Cargo adaptor molecule.

**Figure 5:**
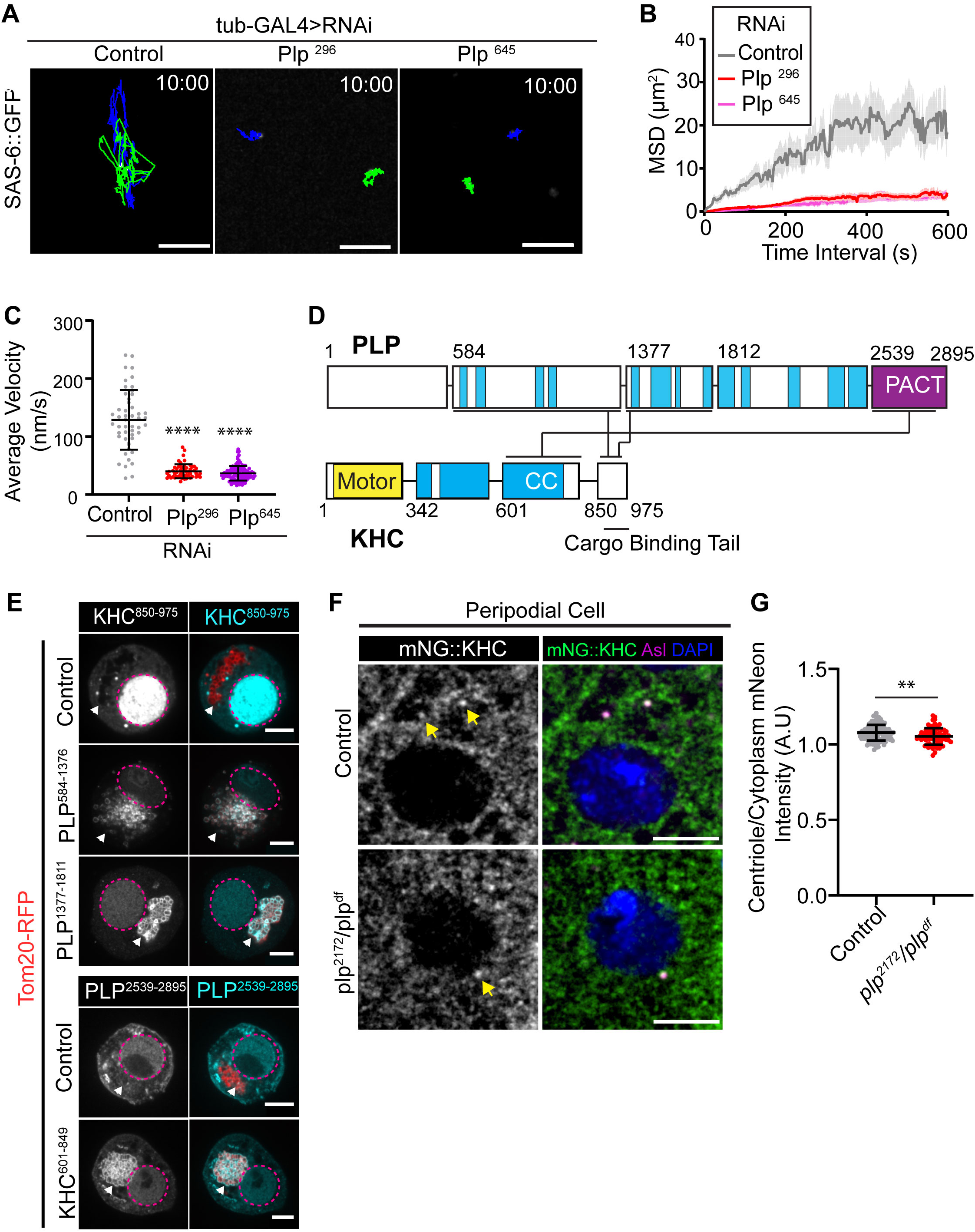
PLP is essential for centriole motility and interacts with KHC. **A)** 10 minute time projections of centriole movement (colored tracks) in PCs following knockdown of PLP with tub-GAL4. Scale bars: 5µm. **B and C)** Quantification of Mean squared displacement (B) and average velocity (C) following PLP knockdown (Control Average velocity = 128.8 nm/s n=49; PlpRNAi296 = 40.19nm/s n =54; PlpRNAi645 = 36.6nm/s n=112, ANOVA: p=<0.0001, Dunnett’s pairwise comparison p = 0.0001 between Ctrl and RNAi conditions, ****). **D)** Diagram showing the three independent interactions (black lines) found through yeast two hybrid screening of PLP and KHC sub-fragments (Fig S4). Blue = Predicted coiled coils. Yellow = KHC motor domain. The PACT domain of PLP is located in the final C-terminal fragment (Purple). **E)** Mitochondrial recruitment assay validating interactions found through Y2H. Prey fragments (Grey/Cyan) accumulate in nucleus (Magenta dashed line). In the presence of a co-transfected bait fragments (red, arrowhead) that is targeted to the mitochondria via a Tom20-RFP tag, a positive interacting fragment (Grey/Cyan) is also recruited to the mitochondria. **F)** Examples showing mNG::KHC (green) localization in control or *plp-* PCs. Yellow arrowheads indicated centriolar localization determined by Asterless staining (magenta). Blue = Nucleus. **G)** Quantification of mNG::KHC localization determined by the mean intensity at the centriole divided by mean cytoplasmic intensity. Mean ± Standard Deviation: Control = 1.078 ±0.05, n=116 centrioles; *plp^2172^/Df* = 1.05±0.06, n=69 centrioles, unpaired t.test: p=0.003, **.

### PLP directly binds KHC

Given that both PLP and Kinesin-1 are required for centriole motility, and that they colocalize on the centriole, we hypothesized that PLP and Kinesin-1 directly interact. To test for direct protein-protein interactions (PPIs), we performed a yeast two hybrid (Y2H) assay using subdivided fragments of PLP and KHC (Figure 5D). The Y2H identified three interactions: PLP^584-1376^-KHC^850-975^, PLP^1377-1811^-KHC^850-975^ and Plp^2539-2895^-KHC^601-849^ (Figure 5D, S4A); the former two interactions are with the KHC cargo binding tail domain. We confirmed these interactions *in vivo* using a mitochondria-targeting assay that utilizes colocalization to test PPIs (Galletta et al., 2014; Schoborg et al., 2015). Using this assay, all three interactions detected by Y2H were confirmed in *Drosophila* cultured cells (Figure 5E). This same assay confirmed the PLP-KHC interaction is conserved between human Pericentrin (PCNT) and the cargo binding domain of human Kinesin-1 heavy chain (KIF5B; Figure S4B,C).

Because PLP physically interacts with the cargo binding domain of KHC (KHC^850-975^), we hypothesized that PLP serves as an adaptor required to anchor Kinesin-1 to the centriole. We therefore compared the localization of mNG::KHC in control and *plp* mutant PCs and found an extremely small (1.9%) decrease in Kinesin levels. Therefore, PLP is not essential for KHC localization to the centriole (Figure 5F,G), thus not a Kinesin-1 adaptor per se, despite being necessary for centriole motility.

If not serving as an adaptor, then PLP might function as an activator, or enhancer of Kinesin-1 motility. To investigate how PLP and KHC may function together, we co-opted an approach previously used to investigate kinesin motility *in vitro* (Kelliher et al., 2018). *Drosophila* S2 cells were transfected with full-length mNG::KHC and another with HALO::PLP^584-1811^. We used PLP^584-1811^ as it encompassed both central fragments of PLP that bound to the cargo domain of KHC (Figure 5D). Whole cell lysates were then extracted from S2 cells and applied to polymerized MTs. Single molecule TIRF microscopy revealed that PLP^584-1811^ and KHC comigrate along MTs (Figure 6 A,B; Movie 11).

**Figure 6:**
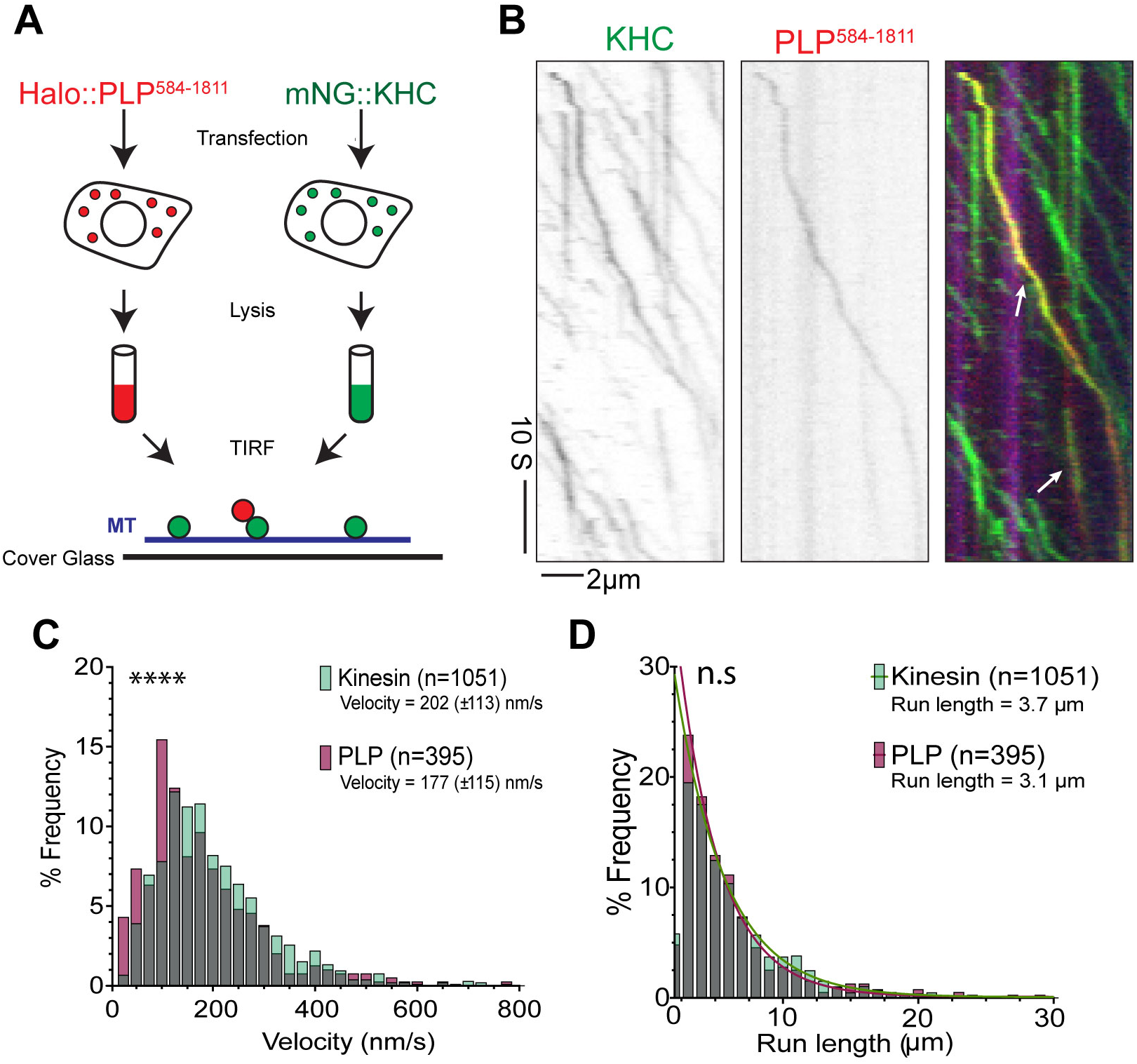
*In vitro* analysis of KHC motility. **A)** Diagram of *in vitro* motility experiment. mNG::KHC (red) and Halo::PLP^584-1811^ (green) were transfected into S2 cells. Cleared lysate was then flowed onto microtubules (blue) for TIRF analysis. **B)** Co-migration of mNG-KHC (green) with Halo-PLP^584-1811^ (red) on HiLyte-647 microtubules (blue). **C)** Velocities of KHC motors and PLP^584-1811^ cargos on microtubules. Kinesin mean velocity = 202 (±113, S.D.) nm/s (n=1051). PLP^584-1811^ mean velocity = 177 (±115, S.D.) nm/s (n=395). Mann-Whitney test P value =<0.0001, ****. **D)** Characteristic run lengths of KHC motors and PLP^584-1811^ cargo. PLP^584-1811^ run length =3.1μm, (n=1051). KHC run length = 3.7μm (n=395). Mann-Whitney test P-value=0.08, ns.

Single-particle tracking showed that Kinesin-1 moved at approximately 200 nm/s, with pauses detected between processive runs. These motile characteristics are consistent with previous studies where full-length Kinesin-1 was shown to undergo frequent pausing whilst moving along MTs, due to the adoption of an autoinhibited state (Friedman and Vale, 1999; Kelliher et al., 2018). Interestingly, our data showed that the dynamics of Kinesin-1 with and without PLP^584-1811^ were similar suggesting that PLP^584-1811^ is not sufficient to activate the Kinesin-1 motor (Figure 6 C,D; Movie 12). This suggests that either another unknown protein is required for Kinesin-1 co-activation with PLP, or that full length PLP is required for activation. Unfortunately we were unable to test these two possibilities as expressing HALO::PLP^FL^ has proven to be exceptionally difficult and another centrosomal interacting partner of KHC is yet to be identified.

### PLP-KHC interaction is required for centriole motility

Although we did not identify the precise mechanism of PLP activation of Kinesin-1, we sought to determine whether the interaction between PLP and Kinesin-1 was required for centriole motility. To test this, we aimed to specifically block the PLP-KHC interaction. This is important as PLP is a large protein that binds multiple proteins to fulfill many functions (Galletta et al., 2016).

To identify *plp* alleles that block Kinesin interaction we took a multifaceted approach (Figure 7A). First, to test significance of the PLP^1377-1811^ – KHC^850-975^ interaction, we used random mutagenesis of PLP^1377-1811^ and screened to identify clones that resulted in loss of Kinesin interaction(Galletta and Rusan, 2015). Screening over 1000 clones yielded one harboring two amino acid substitutions - L1633P and G1699D (*plp^PD^*; Figure S5A,C) that disrupted PLP’s interaction with KHC and Asl while maintaining the integrity of all four other known interactions of this region(Galletta et al., 2016). Second, to test the significance of the PLP^584-1376^ – KHC^850-975^ interaction, we used a series of previously generated deletions and truncations within PLP^584-1376^ because our random mutagenesis approach did not yield a useful allele(Lerit et al., 2015). We found that deletion of amino acids 740-971 (*plp*^Δ740-971^) was sufficient to ablate PLP^584-1376^ interaction with KHC (Figure S5B,C). Finally, we were unable to use random mutagenesis or small deletions to test the PLP^2539-2895^ - KHC^601-849^ interaction, so we completely truncated amino acids 2539-2895 (*plp*^Δ2539-2895^).

**Figure 7:**
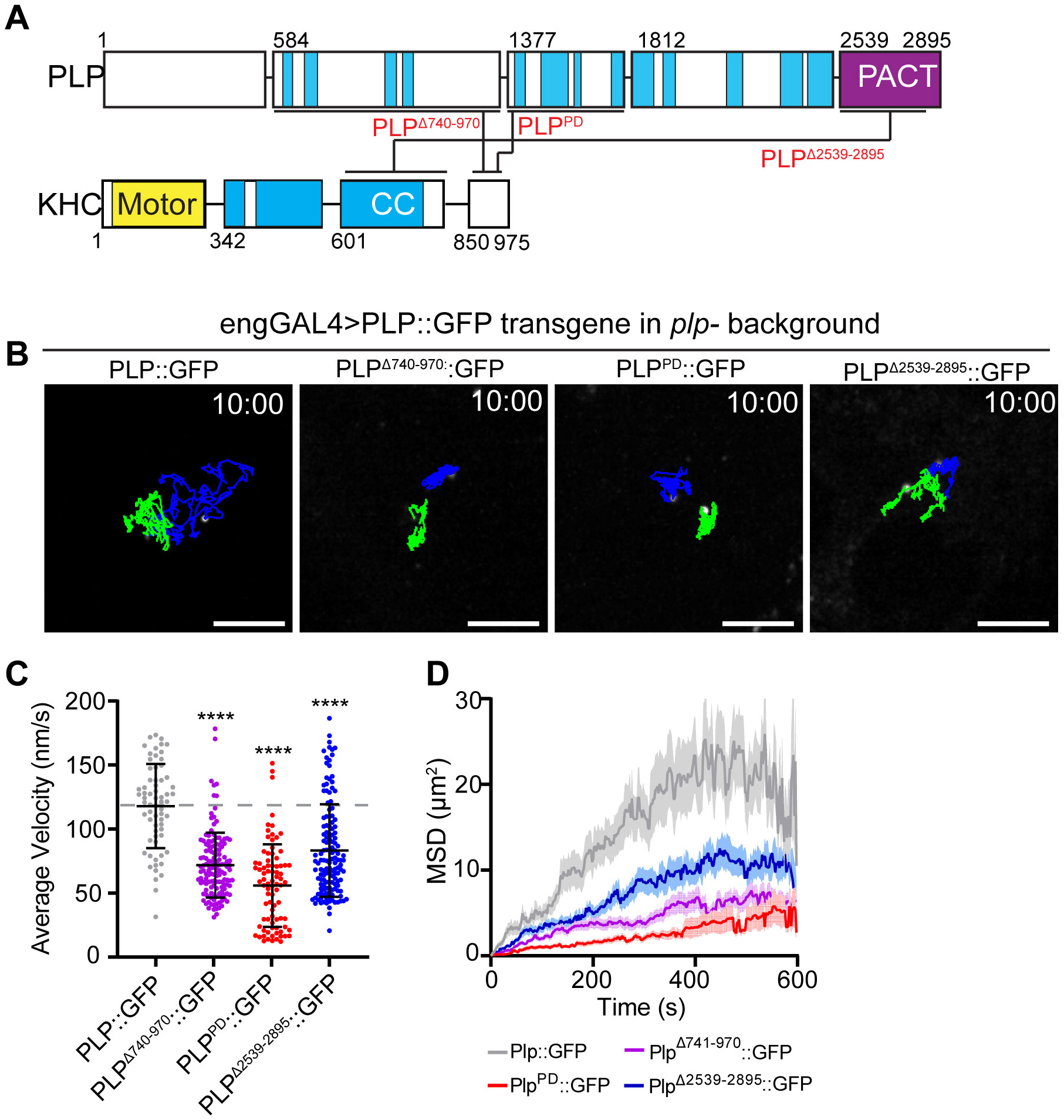
Plp-KHC interaction mutants show reduced centriole motility. **A)** Schematic showing the interactions between PLP and KHC and the corresponding interaction mutations (red text). **B)** 10 minute time projections of centriole movement (colored tracks) in PCs of flies expressing the indicated PLP transgenes in a mutant background (plp2171/Df(3L)Brd15). **C)** Rescue with the mutant transgenes results in a significantly slower instantaneous velocity compared to full length rescue (PLP::GFP: 117±32, n=66. PLP^Δ740-971^::GFP: 71±25, n=83. PLP^PD^::GFP: 55±32, n=138, PLP^Δ2539-2895^::GFP: 83±36 n=139, ANOVA: **** p=<0.0001, Dunnett’s pairwise comparison between PLP::GFP and all other conditions). Data = Mean ± Standard Deviation. **D)** Mean squared displacement shows motility is most effected by mutations (PLP^PD^ and PLP^Δ740-971^) that interfere with interaction between PLP and the cargo binding tail of KHC. Scale bars: 5µm. Time stamp: mm:ss.

To test the physiological relevance of PLP– KHC interaction, we generated transgenic flies expressing each of the three PLP mutant alleles (*plp*^Δ740-971^, *plp^PD^*, *plp*^Δ2539-2895^). Expressing each of these PLP transgenes in the *plp*-mutant background (*plp^2172^/Df*) did not fully rescue centriole motility (Figure 7B-D; Movie 13). The most severe defects were caused by *plp^PD^*, which disrupts the PLP interaction with the cargo binding tail of Kinesin-1. We therefore conclude that PLP interaction with Kinesin-1 is critical for motility.

### PLP and Kinesin-1 are required for PCM asymmetry and proper centriole inheritance

PLP has previously been shown to be involved in three main processes in NB asymmetric division (Figure 8A). Firstly, PLP is required for the timely migration of the mother centrosome away from the apical cortex (Figure 8Aii). Secondly, PLP is required for maintaining the correct number of centrioles/centrosomes in NBs; and thirdly, PLP is required for correct PCM asymmetry between the mother and daughter centrosome (Figure 8B) (Lerit and Rusan, 2013). Whilst Kinesin-1 has previously been implicated in the first of these (Gallaud et al., 2014), its role in the other two processes has not been explored. Unfortunately, our PLP-KHC interaction mutant transgenes (Figure 7) formed large PLP aggregates in NBs, impeding analysis of centriole behavior in this cell type (Figure S5D). We therefore performed a detailed comparison between PLP and KHC loss-of-function to provide insight into which of these three roles of PLP in NBs also involve KHC.

**Figure 8:**
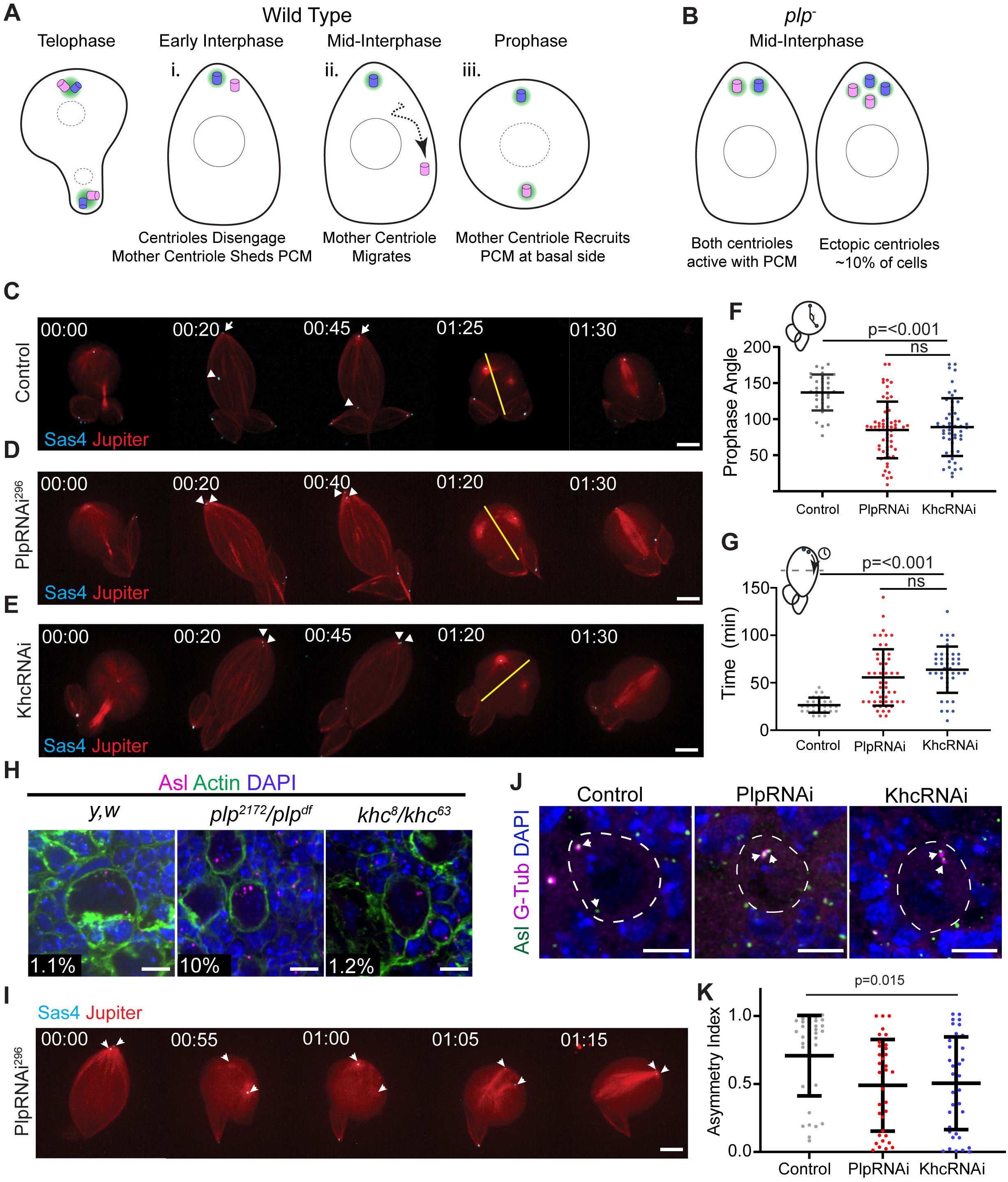
Comparison of PLP and KHC loss of function in NBs. **A)** Schematic of centrosome asymmetry in a wild type NB. i) Mother centriole (Pink) sheds PCM (Green). ii) Mother centriole migrates to the basal side. iii) Mother centriole recruits PCM in the following prophase. **B)** Schematic showing the main phenotypes in *plp*-mutants: Centrioles do not migrate away from the apical domain, mother centriole retains PCM, some NBs inherit supernumerary centrioles. **C, D, E)** Live imaging of NBs expressing the centriole marker mNG::Sas4 (cyan, arrows) and the microtubule marker Jupiter::mCherry (red). The apical daughter centriole is indicated in control NB (arrow), metaphase spindle axis is indicated by yellow line. Unlike controls (C), centrioles do not migrate away from the apical cortex following PLP (D) or KHC (E) knockdown. **F)** The angle between the two centrosomes is significantly reduced at prophase following PLP or KHC knockdown (Control Angle: 137 +/-4.2, PlpRNAi angle: 84.98 +/-4.9, Khc RNAi angle: 88.94+/- 5.5; ANOVA p =<0.0001, Tukey multiple comparison: p=<0.0001 between Control and RNAi conditions), ns = not significant. **G)** The time taken for one centriole to cross the cell midline (equator) is significantly increased following PLP or KHC knockdown compared to controls (Control Separation time: 26.43 +/- 1. PlpRNAi Separation time: 55.65 +/- 4 Khc Separation time: 63.84 +/-3.7; ANOVA p =<0.0001, Tukey multiple comparison: p=0.001 between Control and RNAi conditions). **H)** Representative NBs showing supernumerary centrioles are present in *plp-* (*plp^2171^/df(3L) Brd15*, 10%, n=258 NBs, 4 brains) mutants but not in control (*y,w,* 1.1%, n=229 NBs, 4 brains) or *khc*- (*khc^8^/khc^63^,*1.15%, n=195 NBs, 4brains) mutants. Numbers represent percentage of cells carrying >2 centrioles as determined by Asl puncta (magenta) **I)** Time series showing supernumerary centrioles following *plp-* loss of function arise from a failure of centrosomes to separate in prophase. **J)** Fixed neuroblasts showing gamma-tubulin (magenta) associates with both centrioles (green) following PLP or KHC knockdown. **K)** Quantification of the asymmetric index showing a significant reduction in gamma tubulin asymmetry in PLP or KHC knockdown (Control ASI: 0.7±0.29, n=37. PLPRNAi ASI: 0.49±0.33, n=34. KHCRNAi ASI: 0.5±0.37, n=46; ANOVA: p = <0.0001, Tukey Pairwise comparison: p=<0.0001 between Control and all other conditions). Data = Mean ± Standard Deviation. Scale bars: 5µm.

Firstly, we directly compared the centriole separation phenotype following knockdown of PLP or KHC by live imaging. Analyzing the time required for one of the two centrioles to cross the cells equator in interphase, as well as the angle between the two centrosomes at prophase revealed no significant difference between PLP and KHC knockdown but a significant difference to controls (Figure 8C-G; Movie 14). These results indicate that both PLP and KHC are important for mother centriole motility.

Secondly, we examined centriole number following PLP or KHC loss of function. We confirmed that *plp* mutants have supernumerary centrioles in 10% of NBs; however, *khc* mutants did not (Figure 7H). Importantly, even at low temperature which enhances the *plp* phenotype, *khc* NBs were comparable to controls in centriole number (Figure S6). We concluded that the supernumerary centriole phenotype was due to a function of PLP that is independent of its role in interphase centriole motility. Indeed, live imaging suggests that the centriole number defect in *plp* mutants arises from an inability to separate centrosomes in mitosis, resulting in both centrosomes being inherited by the NB after division (Figure 8I, Movie 15). This is likely due to problems in the Kinesin-5/Dynein dependent centrosome separation pathway that is activated after nuclear envelope breakdown (Tanenbaum and Medema, 2010; Agircan et al., 2014). Loss of *khc* does not appear to effect the Kinesin-5/Dynein pathway in mitosis.

Thirdly, we examined PCM asymmetry on NB centrosomes. If mother centriole motility from the apical to basal side was important for centrosome asymmetry, then we hypothesized that the knockdown of both PLP and Kinesin-1 would perturb the asymmetric localization of PCM on centrioles. Indeed, the asymmetry index (ASI) in NBs expressing PLP or KHC RNAi was significantly reduced compared to controls (Figure 8J,K). In both cases the reduction in asymmetry was due to increased levels of PCM on the mother centriole, rather than loss of PCM from the daughter centriole suggesting that PLP and KHC may be working together for centrosome asymmetry.

Previously, the PCM asymmetry defects observed in *plp* mutants were proposed to be due to its role in inhibiting the localization of the PCM regulator Polo kinase, which functions to promote PCM recruitment (Lerit and Rusan, 2013; Singh et al., 2014). Our finding that KHC knockdown also results in defective PCM asymmetry suggest that Kinesin-1 is also involved in regulating Polo. Consistent with this, live cell imaging of endogenous Polo::GFP localization in isolated NBs revealed that, like PLP, KHC is necessary for Polo asymmetry in early prophase (Figure 9A,B; Movie 16). This raises the possibility that the increase in Polo levels on the mother centriole seen in *plp* mutants may arise via a defect in Kinesin-1 function. How might this occur? One possibility is that Kinesin-1 could directly promote the removal of Polo from the mother centriole. Alternatively, the Kinesin-1 dependent movement of the mother centriole away from the apical microtubule aster, could prevent exposure to Polo activity and thereby prevent precocious PCM recruitment.

**Figure 9:**
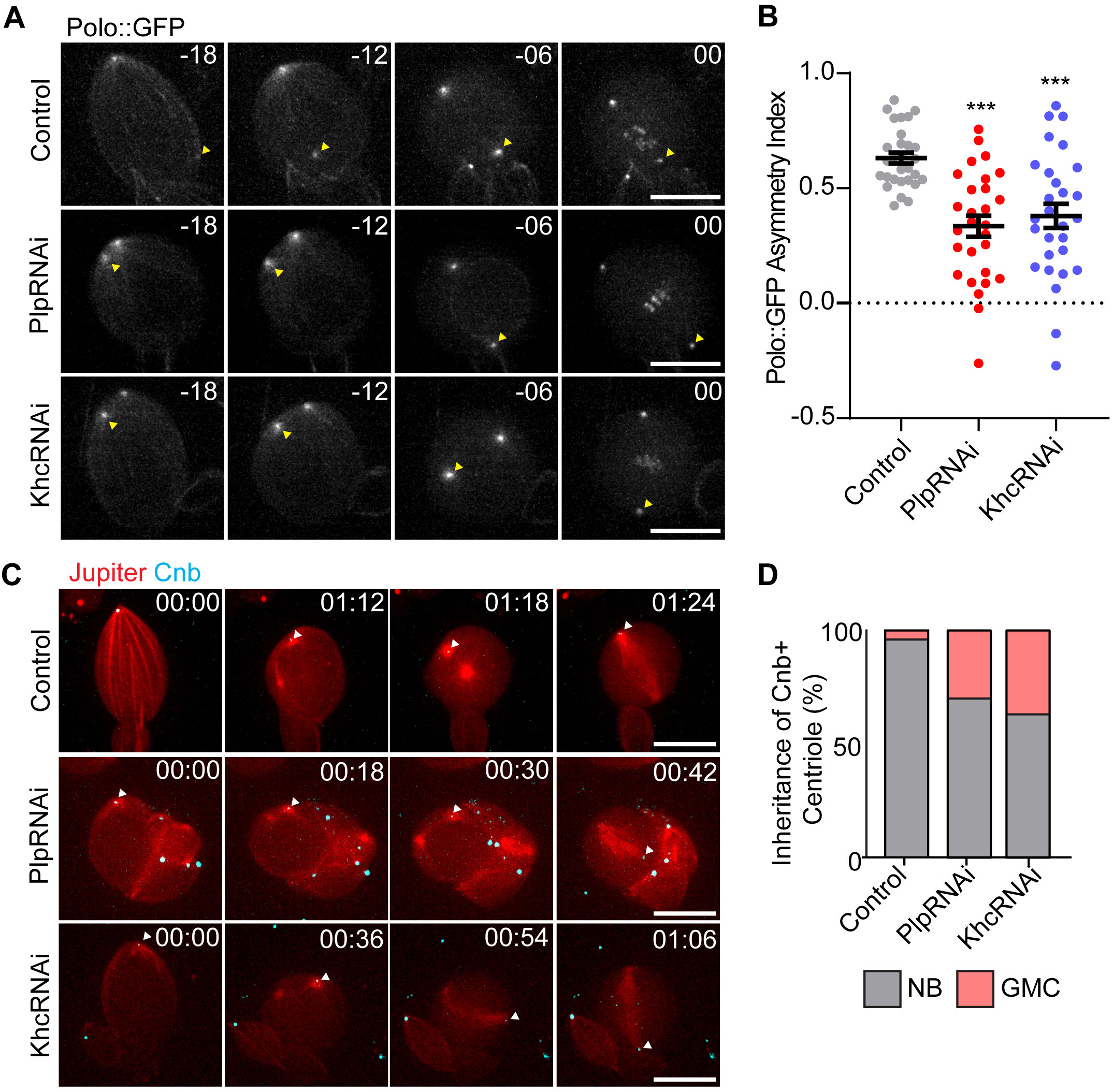
Loss of KHC or PLP causes early Polo exposure and defective age dependent centriole segregation. **A)** Polo::GFP is recruited early to the ganglion mother cell (GMC) inherited centrosome (Yellow arrowhead) in isolated NBs expressing KHC or PoloRNAi **B)** Quantification of Polo asymmetry ∼18minutes before metaphase (Control ASI: 0.63±0.12, n=29. PLPRNAi ASI: 0.33±0.24 n=28, KHCRNAi ASI: 0.37±0.28 n=28, ANOVA: p=<0.0001, Tukey pairwise comparison: Control vs PLPRNAi p = <0.0001, Control vs. KHCRNAi p=0.0002). **C)** Isolated NBs expressing Cnb::GFP (Cyan, arrowhead) to label the daughter centriole. In NBs also expressing KHC or PLP RNAi, the daughter centriole is more frequently segregated into the GMC. D) Quantification of Cnb::GFP+ centriole inheritance (% Cnb+ Centriole Inherited by NB: Ctrl: 96%, PLPRNAi: 74%, KHCRNAi: 63%, 40 NBs were imaged per condition). Scale bars: 10µm.

NBs consistently retain the daughter centriole during asymmetric cell division. We hypothesized that given PCM asymmetry is perturbed prior to separating during interphase, that the age dependent segregation of centrioles would be disrupted. To examine this, we performed live cell imaging of neuroblasts expressing GFP tagged Centrobin (Cnb), a marker for the daughter centriole (Januschke et al., 2011, 2013). Observing the segregation of the Cnb+ centriole revealed that in the absence of PLP or KHC the daughter centriole was more likely to be segregated to the GMC compared to controls (Figure 9C,D; Movie 17). We conclude that KHC and PLP are similarly important to generate correct PCM asymmetry which facilitates daughter centriole retention in NBs.

## Discussion

The movement of centrioles is critical for an array of cellular processes. Most of the work in this research area has focused on the separation of active centrosomes (not inactive centrioles), which move by indirect transport using motors anchored to the cortex or anchored to other MTs (sliding; Tanenbaum and Medema, 2010; Agircan et al., 2014). In this work we have leveraged *Drosophila* to understand how centrioles can move when they are not functioning as a MTOC. In multiple cell types, we show that centrioles move on MTs as cargo, independently of MT nucleation (Figure 3). Several of our results support a model whereby centrioles are cargo for the motor protein Kinesin-1. Firstly, the knockdown of Kinesin-1 components inhibits motility, independent of KHC’s role in MT sliding (Figure 4). Secondly, KHC localizes to the outer centriole edge. Thirdly, KHC interacts with PLP, and this interaction is necessary for centriole motility (Figures 5-7). Interestingly, PLP is not necessary for the localization of KHC to the centriole, suggesting that it is likely contributing to motility by somehow activating Kinesin-1 the motor rather than merely anchoring the motor to the centriole.

Although PLP^584-1811^ was able to co-migrate with KHC on MTs *in vitro*, it alone was not sufficient to enhance the basal activity of the Kinesin-1 motor *in vitro* (Figure 6). It is not uncommon that Kinesin-1 requires multiple interactors for full activation. One example of this is the scaffolding protein JNK-interacting protein 1 (Jip-1) which is insufficient alone for Kinesin activation but instead co-operates with secondary factors for activation (Blasius et al., 2007; Hammond et al., 2008; Sun et al., 2011).

Interestingly, this complexity of motor activation is likely a mechanism by which cells can control the directionality of intracellular transport in cases where a single cargo can bind multiple types of motors (Fu and Holzbaur, 2014). For example, many adaptor molecules can interact with both plus-end directed Kinesin-1 and minus-end directed dynein, which interestingly has been implicated in the movement of centrioles into the oocyte during *Drosophila* oogenesis (Grieder et al., 2000; Bolvar et al., 2001). Moreover, in mammalian cells Dynein intermediate chain interacts with Pericentrin (Purohit et al., 1999; Young et al., 2000; Sepulveda et al., 2018). Given that we were searching for a plus-end directed motor for centriole transport in NBs, we have not investigated Dynein.

Why are centrioles motile? During asymmetric cell division of NBs the mother centriole sheds PCM following cytokinesis and migrates away from the daughter which recruits PCM and becomes an MTOC. We show that the motile mother centriole is associated with the MT network (Figure 3B) and appears to move along it directionally (Figure 3D). Kinesin-1 and PLP directly interact (Figure 5) and are required for the timely separation of centrioles prior to prophase (Figure 7A; Lerit and Rusan, 2013; Gallaud et al., 2014). Therefore, we propose that PLP and Kinesin-1 are functioning together for centriole separation in NBs.

What is the consequence of delayed centriole separation? *plp^-^* mutants were previously shown to accumulate ectopic centrioles (Lerit and Rusan, 2013); however, we did not observe ectopic centriole formation in *khc* mutant NBs. This suggests that the supernumerary centrioles in *plp^-^* mutants is not entirely caused by defective interphase centriole motility. Instead, it appears to also result from defective centrosome separation in prophase and beyond, likely due to the role PLP has in organizing mitotic PCM and MTs (Galletta et al., 2014; Lerit et al., 2015; Varadarajan and Rusan, 2018).

Another consequence of impaired motility is defective centrosome asymmetry. Normally through interphase, PCM is restricted to the daughter centriole, allowing the mother to effectively migrate away (Figure 1; Rebollo et al., 2007; Rusan and Peifer, 2007; Conduit and Raff, 2010; Lerit and Rusan, 2013; Singh et al., 2014). Interestingly, both KHC and PLP knockdown caused defective centrosome asymmetry. How might centriole movement contribute to the asymmetric recruitment of retention of PCM? Polo is a critical regulator of PCM recruitment. Polo localizes to both the centriole and the MT network, where it has been proposed to move toward the apical centrosome (Ramdas Nair et al., 2016). We show that in the absence of centriole motility, Polo::GFP is present on both mother and daughter centrioles in prophase (Figure 9A). We propose that this is due to the high local concentration of Polo at the apical MT aster (Ramdas Nair et al., 2016); when the mother centriole is unable to migrate away, and maintains its position at the apical pole, it would be exposed to these higher levels of Polo. It is important to note that we cannot formally rule out that PLP and Kinesin-1 are functioning together to antagonize Polo independently of centriole motility.

Age dependent segregation is a conserved phenomenon (Chen and Yamashita, 2021). Why does the age dependent segregation of centrosomes matter? This is unclear. Reports have shown that fate determinants (Ramat et al., 2017; Tozer et al., 2017), damaged proteins (Fuentealba et al., 2008) and foreign DNA(Wang et al., 2016) are all associated with centrosomes of a determined age. Without an experimental system to trigger inverse age segregation in every cell cycle it will be difficult to determine the true functionality of age dependent centriole segregation.

Is Kinesin-1 mediated centriole motility conserved? In human cells, centrosomes have been observed to be motile at the end of mitosis, when the centrosomes reorient themselves towards the midbody (Piel et al., 2001). Interestingly, recent work has demonstrated that this centrosome motility involves the rab11 dependent localization of PCNT to the centrosomes (Krishnan et al., 2021). It is unclear whether Kinesin-1 is involved in this process, but Kinesin does localize to centrosomes (Neighbors et al., 1988) and interacts with PCNT (Figure S4).

In summary we have dissected a novel mechanism for centriolar transport where PLP functions as a Kinesin-1 activator to facilitate direct MT transport of centrioles. Importantly, we have shown that this process is important for centriole separation in asymmetric cell division. Future studies will be required to uncover how Kinesin-1 is activated at the centriole to ensure proper centriole motility.

## Methods

### Plasmids and Molecular Cloning

Sequences were amplified from cDNA clones by PCR and inserted into the pEntr gateway vector using the pEntr D-Topo kit (Life Technologies). Gateway reactions were used to recombine the cDNA into the destination vectors. ANW: Actin promoter with N-term mNeon tag. PAT20RW: Actin promoter with N-term TOM20 mitochondrial localization domain and TagRFP. PPWG: UAS promoter with C-term GFP. pDEST-pGADT7: Y2H bait plasmid, GAL4 binding domain. pDEST-pGBKT7: Y2H prey plasmid with GAL4 activation domain. PCNT and KIF5B fragments were generated using DNA synthesis (Twist Biosciences). The following published plasmids were used: pEntrPLP^1-583^, pEntr-PLP^584-1375^, pEntr-PLP^1376-1810^, pEntr-PLP^1811-2537^, pEntr-PLP^2538-2895^, pEntr-PLP^583-1810^ and pEntr-Sas6 (Galletta et al., 2016); pEntr-PLP (Galletta et al., 2014); F-Tractin::mCherry (Liu et al., 2021). KHC fragments were amplified with the following primers: KHC^1-341^ Fwd: 5’-CACCATGTCCGCGGAACGAGAGATTCC-3’, Rev: 5’ - CTCGTTAACGCAGACCACGTTCTTCACTGTCTTG-3’; Khc^342-600^ Fwd: 5’ - CACCATGGAGCTTACTGCCGAGGAATGGAAG-3’, Rev: 5’ - AGCCAGAGCACTCATCTTAAGGTCGATGCTGG-3’; KHC^601-849^ Fwd: 5’ - CACCATGGGCACGGATGCCAGCAAG-3’, Rev: 5’ -CGCGAGTGATCCACCGTCCTCCTC-3’; KHC^850-975^ Fwd: 5’ -CACCATGCAGAAACAGAAGATTTCCTTCTTGGAGAACAACC-3’, Rev: 5’– CGAGTTGACAGGATTAACCTGGGCCAGC-3’.

### Fly Stocks

Flies were maintained on Bloomington Recipe Fly Food (LabExpress). Crosses were performed at 25°C unless otherwise stated in the text. Transgenic Flies were generated using standard P-element transformation methods (Bestgene). The following stocks were used: *ubi-gfp::sas6* (this study), *sas4::neon*(Galletta et al., 2020), *UAS-lifeact::rfp* (BDSC #58362), *UAS-jupiter::mcherry* (Gift from C. Cabernard), *worniu-GAL4* (BDSC #56553), *AGIR-GAL4* (BDSC #6773), *tubulin-GAL4* (BDSC #5138), *ubi-tubulin::gfp (Gift from T. Avidor-Reiss)*, *ubi-sas6::mcherry*(Rogers et al., 2008), *ubi-cbn::gfp*(Lerit and Rusan, 2013), *polo::gfp*(Ramdas Nair et al., 2016) (Gift from C. Cabernard), *UAS-plpRNAi^296^* (VDRC #101296), *UAS-plpRNAi^645^* (VDRC #101645), *UAS-khcRNAi* (VDRC#44338), *UAS-klcRNAi* (BDSC #42597), *UAS-ensRNAi* (BDSC #40825), *UAS-mcherryRNAi* (BDSC #35785), *UAS-luciferaseRNAi* (BDSC #31603), *UAS-eb1::gfp*(Swider et al., 2019), *UAS-plp::gfp*, *UAS-plp^PD^::gfp* (This Study), *UAS-plp^Δ741-970^::gfp* (This Study), *plp2172* (BDSC #12089), *Df(3L)Brd15* (BDSC #5354), *khc8* (BDSC#1607), *khc63*(Djagaeva et al., 2012) (Gift from B. Saxton), *khc^mut.A^*(BDSC: 79036), *ubi-mNG::KHC* (This study).

### Immunofluorescence and fixed cell microscopy

Larvae were selected at the third instar stage and dissected in PBS before being fixed in 4% formaldehyde at room temperature for 20 minutes. Fixed samples were washed and then blocked in PBS+0.5%Triton-x100 (PBST) + 5% normal goat serum. Samples were then incubated in PBST+primary antibody overnight at 4°C. S2 Cells were cultured in serum free SF900 II media and plated onto coverslips precoated with Concanalavin-A. After being allowed to adhere for 30 minutes SF900 media was removed and replaced with 4% formaldehyde in PBS for 20 minutes. Staining was then performed as with tissue but with 0.1% Triton-x100 rather than 0.5%. The following antibodies were used: anti-Cnn (Galletta et al., 2016) (rabbit, 1:10000), anti-asterless (Klebba et al., 2013) (Guinea Pig, 1:30000), anti-Plp (Rogers et al., 2008) (rabbit, 1:12000), anti-gamma-tubulin (Mouse, 1:500, GTU-88, Sigma-Aldrich), anti-tubulin (Mouse, 1:200, Sigma). Samples were washed in PBST and then incubated in secondary antibody (1:500, Life Technologies) for 2 hours at room temperature. Following secondary antibody incubation, samples were washed in PBS before being mounted in a small drop of vectashield within a reinforcement label used as a spacer. All slides were mounted underneath a #1.5 coverslip. Phalloidin-Alexa488 (5:200, Life Technologies) was added for 20 minutes after the secondary antibody step.

Standard resolution imaging was performed using a Yokogawa Csu-W-1 spinning disk mounted on a NIkon Ti-2 Eclipse equipped with a Prime BSI CMOS camera and a 100x Silicone immersion objective (N.A 1.4). Microscope was controlled using Elements (Nikon).

Structured Illumination Microscopy was performed using an OMX4 (GE healthcare) using immersion oil RI 1.516. Images were reconstructed using SoftWoRx (GE Healthcare). Stimulated Emission Depletion (STED) images were acquired using a Leica SP8 3X STED microscope, a white-light laser for fluorescence excitation (470–670nm), a Leica HyD SMD time-gated photomultiplier tube and a Leica 100x (NA 1.4) STED White objective (Leica Microsystems, Inc.). ATTO 647 and Alexa 594 were excited at 647nm and 575nm with fluorescence emission collected over a bandwidth of 658–755nm and 583–700nm respectively. A 25 slice z-stack for both colors was acquired with a pinhole size of 0.7 airy unit (A.U.), a scan speed of 600Hz, a pixel format of 1024×1024 (pixel size, 20nm), an interslice distance of 0.16μm, four line averages, and time gating on the HyD SMD set to a range of 0.7–6.5ns. HyD SMD gains were set to 100% and 150% for ATTO 647 and Alexa Fluor 594 respectively.

STED depletion was accomplished for both labels at 775-nm (pulsed at 80MHz) at powers of approximately 192mW at the back aperture for ATTO 647-labeled microtubules (40% full laser power) and 99mW for Alexa Fluor 594-labeled centrioles (20% full laser power). STED images were deconvolved using the software Huygens Professional (v.19.1, Scientific Volume Imaging) using an idealized point spread functions and the classic maximum-likelihood estimation deconvolution algorithm.

### Sample preparation and Live cell microscopy

Wing Discs were dissected from third instar larvae in Schneider’s medium supplemented with glucose (1g/l). They were mounted in a drop of the same media on a 50mm gas permeable dish (Lumox, Sarstedt). A drop of halocarbon oil (sigma) was added around the drop of media and a 22x22mm coverslip (#1.5, Thermofisher) was gently lowered on top. Centriole movement was imaged by mounting the wing discs with the peripodial membrane facing the coverslip.

For live imaging of neuroblasts, brains were dissected in collagenase buffer and incubated in collagenase for 20 minutes. Brains were then washed in Schneiders medium supplemented with FCS, Fly Extract (DGRC) and insulin (sigma). They were then dissociated as previous described(Pampalona et al., 2015) and plated onto a 35mm glass bottom dish (Fluorodish, World Precision instruments) precoated with Poly-L-Lysine (Sigma). Cells were allowed to adhere for 45 minutes prior to imaging.

S2 Cells were plated directly onto 35mm Glass bottom dishes coated with Concanalavin-A and allowed to adhere for 30 minutes. Cells were then washed with Schneider’s medium to remove SF900 II.

Imaging was performed on an inverted microscope (Eclipse Ti, Nikon) fitted with a Csu-22 spinning disk confocal head (Yokogawa) and a sCMOS camera (Orca Flash 4, Hammamatsu). Acquisition was set up through the Metamorph software (Molecular Devices). Unless otherwise stated in the figure legend. Wing Disc images were collected at 2s time intervals for 10 minutes total acquisition of a single z slice. Focus was adjusted manually during acquisition when necessary. Neuroblast movies were acquired by taking a 10-12μm volume every 180s using 800nm z-intervals. S2 cell movies were performed by acquiring a 2.4μm volume every 2s using 800nm intervals.

### Particle Tracking

Movies were processed by subtracting the background and then applying a 1px gaussian filter. Tracking was performed using a modified program in IDL (Harris Geospatial) based on a previous particle tracking pipeline (Crocker and Grier, 1996) which can be found here for IDL and other implementations (http://www.physics.emory.edu/faculty/weeks//idl/).

### Drug Treatments

To depolymerize microtubules, wing discs were dissected and placed in Schneider’s medium in an ice/ethanol slurry for 1 hour. Wing discs were then transferred to room temperature media and allowed to recover in the presence of DMSO or 50µM Colcemid (Tocris Bioscience). To block microtubule polymerization wing discs were incubated in 200µM Colchicine at room temperature for 20 minutes before imaging. To block actin polymerization, wing discs were incubated in 10µM Latrunculin-A (Sigma) for 20 minutes before imaging. For drug experiments imaging was performed in the presence of the inhibitor.

### Cell extract preparation

Cell lysate-based motility assays were performed as described previously (Ayloo and Holzbaur, 2015). Briefly, *Drosophila S2* cells were transfected with mNeonGreen-tagged kinesin-1 full-length heavy chain (KHC) and Halo-tagged PLP^584-1811^ (Effectene, QIAGEN). The lysates were harvested for TIRF motility assays at ∼48 h post-transfection. Prior to extraction, HaloTag was labeled with tetramethylrhodamine (TMR) by incubating the transfected cells with 2.5 mM HaloLigand-TMR (Promega) for 15 min. Unbound ligand was washed out with PBS, and the cells were lysed in buffer containing 120 mM NaCl, 0.1% Triton, 1 mM ATP, 1 mM EGTA and 40 mM HEPES (pH 7.4) supplemented with protease inhibitor cocktail (Roche). The cell extract was centrifuged at 100,000 g for 10 min at 4 °C and the supernatant was kept on ice until use.

### TIRF motility assays

Unlabeled (T240), HiLyte-647 (TL670M) and Biotin (T333P) labelled porcine tubulins were purchased from Cytoskeleton. Tubulins were resuspended according to the manufacturer’s instructions and mixed at ratio of 50:1 (Unlabeled: Biotin) or 50:1:1 (Unlabeled: Biotin:Hilyte-647), with a total tubulin concentration of 10 mg/ml, in ice cold BRB80 (80 mM K-PIPES, 2 mM MgCl2, 1 mM EGTA (pH 6.8 with KOH)) with 1 mM GTP. Polymerization was performed by adding warmed (37°C) BRB80 + 1 mM GTP + 10% glycerol and incubating at 37°C for 30 minutes. For stabilization, Taxol was added stepwise (0.5 µM, 5 µM and 50 µM after addition) with 10 minutes incubation at 37°C for each step.

10µl flow chambers with biotin-PEG functionalized coverslips were constructed as described previously (Tripathi et al., 2021). Chambers were washed with BRB80 (3x10 µl) followed by 5 mg/ml BSA + 5 mg/ml casein in BRB80 (3x10 µl). The final chamber volume was incubated in the chamber for 1 minute. 2 mg/ml Neutravidin (1x10 µl) in BRB80 was incubated in the chamber for 1 minute and then washed with BRB80 + 20 µM Taxol (3x10 µl). 0.5 µM microtubules in BRB80 + 20 µM Taxol + 50 mM DTT were added and incubated for 1-3 minutes to allow for sufficient surface attachment. The chamber was washed with BRB80 + 20 µM Taxol + 50 mM DTT (3x10 µl). KHC, PLP^584-1811^ or KHC + PLP^584-1811^ cell lysates were diluted into assay buffer (BRB80 + 20 µM Taxol, 50 mM DTT, 1 mM ATP, 100 μg/ml glucose oxidase, 40 μg/ml catalase, 2.5 mg/ml glucose) and flowed into the chamber for imaging (3x10 µl). Optimal dilutions and mixing ratios were determined empirically based on the quantity of motility observed.

Movies were collected on an inverted Nikon Eclipse Ti-E microscope with H-TIRF module attachment, CFI60 Apochromat TIRF 100× Oil Immersion Objective Lens (N.A. 1.49, W.D. 0.12 mm, F.O.V 22 mm) and an EM-CCD camera (Andor iXon Ultra 888 EMCCD, 1024 × 1024 array, 13 μm pixel). For movie 12, a maximum intensity projection of the KHC channel was performed to highlight the position of the microtubule track (depicted in blue). Particles were tracked using the FIJI plugin Trackmate (Tinevez et al., 2017) and plotting/analysis was performed in Graphpad Prism. Pauses between movements were included in this analysis. Data from three separate movies were combined to produce each of the KHC and PLP^584-1811^ datasets shown in figure 5. All quantifications relate to the mixture of both proteins.

### Yeast-2-Hybrid Screening

Yeast 2 Hybrid experiments were performed using a previously described modified version of the Matchmaker Gold system (Galletta and Rusan, 2015; Galletta et al., 2016). Constructs listed were recombined into bait and prey plasmids through gateway recombination. Bait plasmids were transformed into the Y2H Gold strain and Prey plasmids into Y187. Strains were individually mated in 2x Yeast extract Peptone Adenine dextrose. Plates were incubated at 30°C before being plated on DDO (SD-Leu-Trp) plates to select diploids carrying both prey and bait plasmids. Colonies were then replica plated onto DDO, QDO (Sd-Ade-His-Trp), DDOXA (DDO+Auereobasidin A + X - α -Gal) for selection. Plates were scored based on the presence or absence of robust blue yeast growth on DDOXA plates.

### Identification of Plp-KHC interaction mutations

Three fragments of Plp (Plp^584-1376^, Plp^1377-1811^, Plp^2539-2895^) were found to interact with KHC. To identify mutants that disrupted these interactions we first used a method previously described(Galletta et al., 2016). To induce random mutagenesis, Plp fragments were amplified by low fidelity PCR, a result of limiting dATP concentration in the PCR reaction (0.06mM dATP, 0.25 mM dCTP, dGTP and dTTP). This will induce a mutation approximately every 250bp. Primers used: Fwd: 5’ -CGGAATTAGCTTGGCTGC-3’, Rev: 5’ -TAATACGACTCACTATAGGGCG-3’. Amplification products were co-transformed into Y2H Gold with linearized pGBKT7. Approx. 2000 clones were isolated of each mutagenesis reaction and arrayed in 96 well plates before being mated with the corresponding KHC fragment (in pGADT7, Y187 strain). The Array was then screened on plates for loss interaction (no growth on QDOXA and DDOXA plates). Clones that did not interact were isolated from the parent array and retested against previously identified interactors of Plp(Galletta et al., 2016). Using this approach, we were successful in identifying a Plp^1377-1811^ clone that retained interaction with most other identified interactors, but not KHC or Asl (Supp fig. 5A). Sequencing revealed this clone carried mutations resulting in two substitutions (L1633P and G1699D). We referred to this mutant as PLP^PD^. The yeast mutagenesis approach did not yield useable mutations in either _PLP_584-1376 _or PLP_2539-2895.

The next approach was to screen a series of deletions within these two fragments for loss of KHC interaction. Previously a series of deletions had been generated within PLP^584-1376^ (Lerit et al., 2015). We found that a deletion corresponding to amino acids 741-970 inhibited interaction with KHC and the C-terminus of PLP without effecting other known interactors (Supp Fig 5B). We were unable to identify any deletions within PLP^2539-2895^ that inhibited Kinesin interaction.

Insertion of the Plp^Δ741-970^ into the full length PLP transgene was performed by amplifying the region downstream and upstream of the deletion and then assembling them using Gibson assembly into the pEntr plasmid. Primers used were as follows: F1 Fwd 5’ – GCGGGAAAAACACATTCCTCCCTACCTCCA-3’, F1 Rev 5’ - CAGAGTTTTAGACTCATCCAAGGAGAGGGA-3’, F2 Fwd 5’-TCCCTCTCCTTGGATGAGTCTAAAACTCTG-3’, F2 rev 5’ – TGGAGGTAGGGAGGAATGTGTTTTTCCCGC -3’.

Insertion of PLP^PD^ into the full length PLP transgene was performed by Gibson assembly of two fragments using the following primers: F1, Fwd 5’ -GGTTGCGTGGAGCTTCAACATGAGC-3’, Rev, 5’ -GATGCATTTCCCGCATGCTCTTGAAGATC-3’. F2, Fwd 5’ - GCGGGAAAAACACATTCCTCCCTACCTCCAGATCTTCAAGAGCATGCGGGAAATGCATCA-3’, Rev, 5’ –AGTCTGTTCCTCCATACGACCCTGCAGCGTATCCCGCTCATGTTGAAGCTCCACGCAACC-3’.

The cloning of PLP^Δ2539-2895^ was previously described(Lerit et al., 2015).

## Acknowledgements

We would like to thank the Xufeng Wu and the NHLBI Light microscopy core facility for technical assistance. We would like to thank all members of the Rusan lab for discussion. We thank Ryan O’Neill, Chaitali Khan, Alexander Kelly and Greg Rogers for critically reading the manuscript. We would like to thank Sam Smith for discussions about single molecule analysis of PLP. Finally, we thank Clemens Cabernard, Bill Saxton, Jill WiIdonger and Tomer Avidor-Reiss for kindly sharing reagents. This work is supported by the Division of Intramural Research at the National Institutes of Health/National Heart, Lung, and Blood Institute (1ZIAHL006126 to N.M.R. and ZIAACTHL006049 to J.R.S.)

## Author contributions

M.R.H, R.L, N.B., Z.T.S., B.J.G, C.J.F performed the experiments and analyzed the data. C.C. provided critical assistance in particle tracking and super resolution microscopy. M.R.H. and C.J.F designed and cloned all constructs. M.R.H and N.M.R wrote the manuscript. J.R.S and N.M.R. supervised and funded the project.

---

**Supplementary Figure 1:**
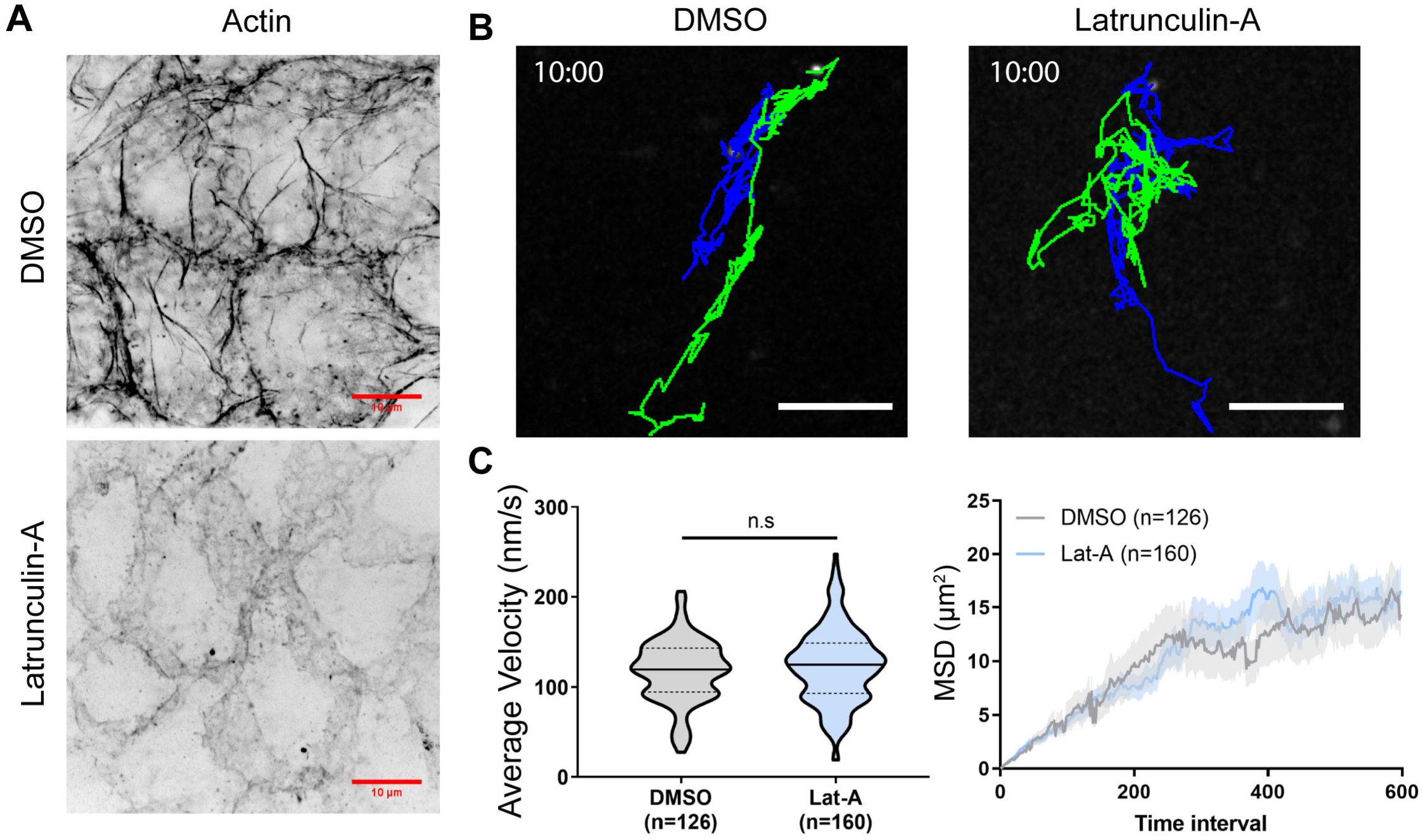
Centriole motility is independent of the Actin network. **A**) Z-stack projection of example PC’s labelled with Phalloidin showing that 10µM Latrunculin-A treatment destroys the Actin network. Scale bars: 10μm. **B)** Tracks showing the movement of centrioles over a 10 minute period. Latrunculin-A treatment does not inhibit centriole motility in peripodial cells. Scale bars: 5μm. **C)** Quantification of average velocity. (DMSO: 117.6 nms ± 34, n= 126, Lat-A: 123.5 nms ± 41, n= 140. T.Test: p=0.19). **D)** Mean squared displacement is not affected by Latrunculin A treatment. Data: Mean ± Standard Deviation.

**Supplementary Figure 2:**
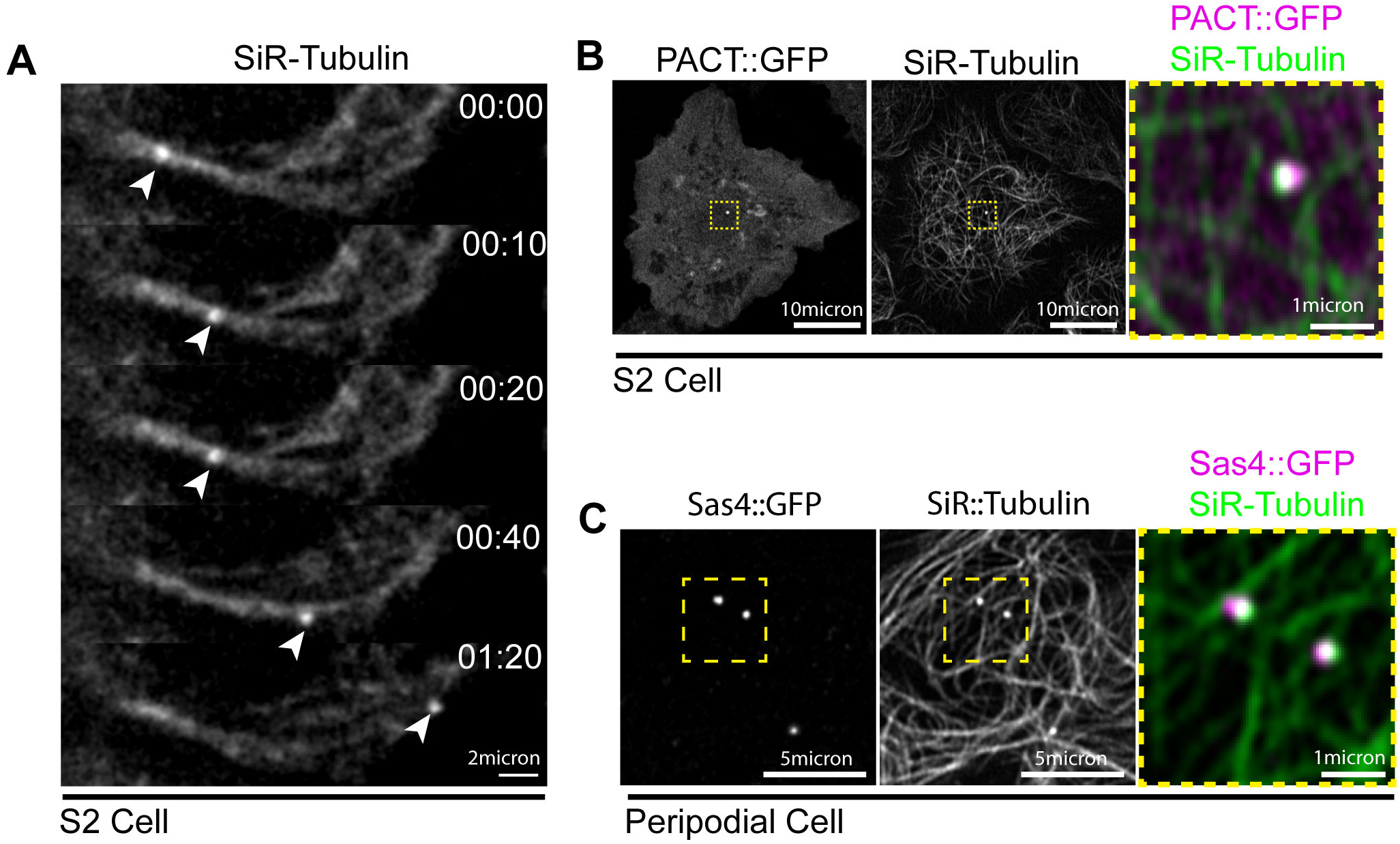
SiR-tubulin labels centrioles and microtubules. **A)** Timelapse series of an S2 cell labelled with SiR-Tubulin. Arrowhead denotes brighter spot corresponding to the centriole moving along the microtubule network. **B)** Projection of a live S2 cell transfected with PACT::GFP to label the centriole (Magenta). Centriolar signal is coincident with bright accumulation of SiR-Tubulin (Green). **C)** Live PC expressing Sas4::GFP (centriole, magenta) and labelled with SiR tubulin (Green). SiR tubulin accumulation corresponds to Sas4 positive centrioles.

**Supplementary Figure 3:**
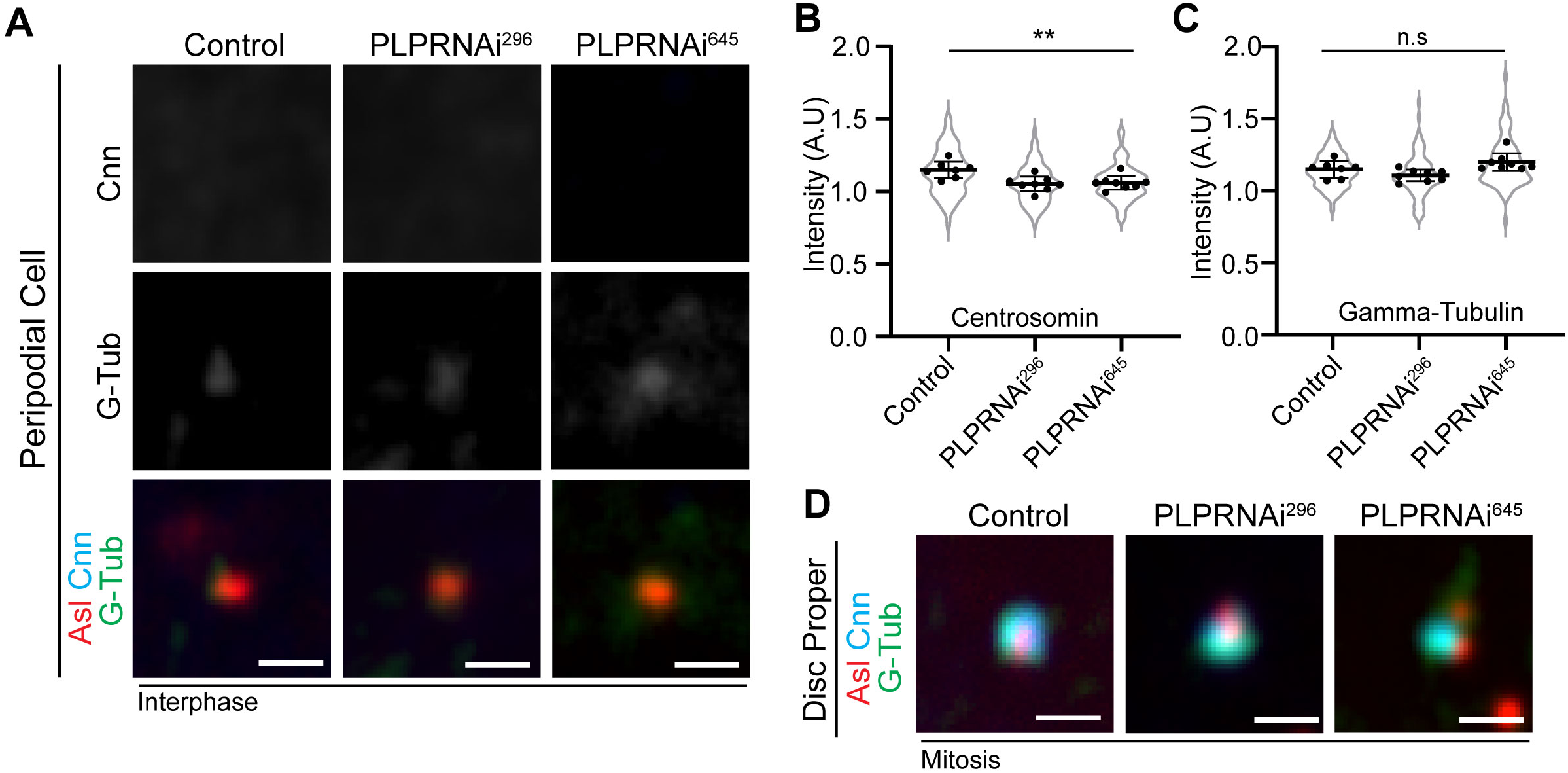
Plp knockdown does not cause precocious centriole activation in PCs. **A)** Fixed PCs stained for Asl (red), Cnn (cyan) and Gamma-tubulin (red) shows that PCM does not accumulate in PLPRNAi expressing PCs. **B)** Quantification of Centrosomin intensity relative to cytoplasm (Control: 1.16±0.05 n= 7 wing discs, 67 centrioles; PLPRNAi^296^: 1.1±0.05, n= 8 wing discs, 91 centrioles; PLPRNAi^645^: 1.1±0.05, n= 8 wing discs, 86 centrioles. ANOVA: p= 0.003. Dunnett’s multiple comparison: Control vs PLPRNAi^296^: p = 0.0035, Control vs PLPRNAi^645^: p = 0.0065). **C)** Quantification of Gamma Tubulin intensity relative to cytoplasm (Control: 1.15±0.05 n= 7 wing discs, 67 centrioles; PLPRNAi^296^: 1.1±0.04, n= 8 wing discs, 91 centrioles; PLPRNAi^645^: 1.2±0.06, n= 8 wing discs, 86 centrioles. ANOVA: p= 0.01, multi-comparison test showed no significance between Control and RNAi groups. **D)** Fixed mitotic wing disc cells stained for Asl (red), Cnn (cyan) and Gamma-tubulin (red). Plp knockdown does not prevent PCM accumulation in mitosis. But PCM can appear disorganized. Scale bars: 1µm.

**Supplementary Figure 4:**
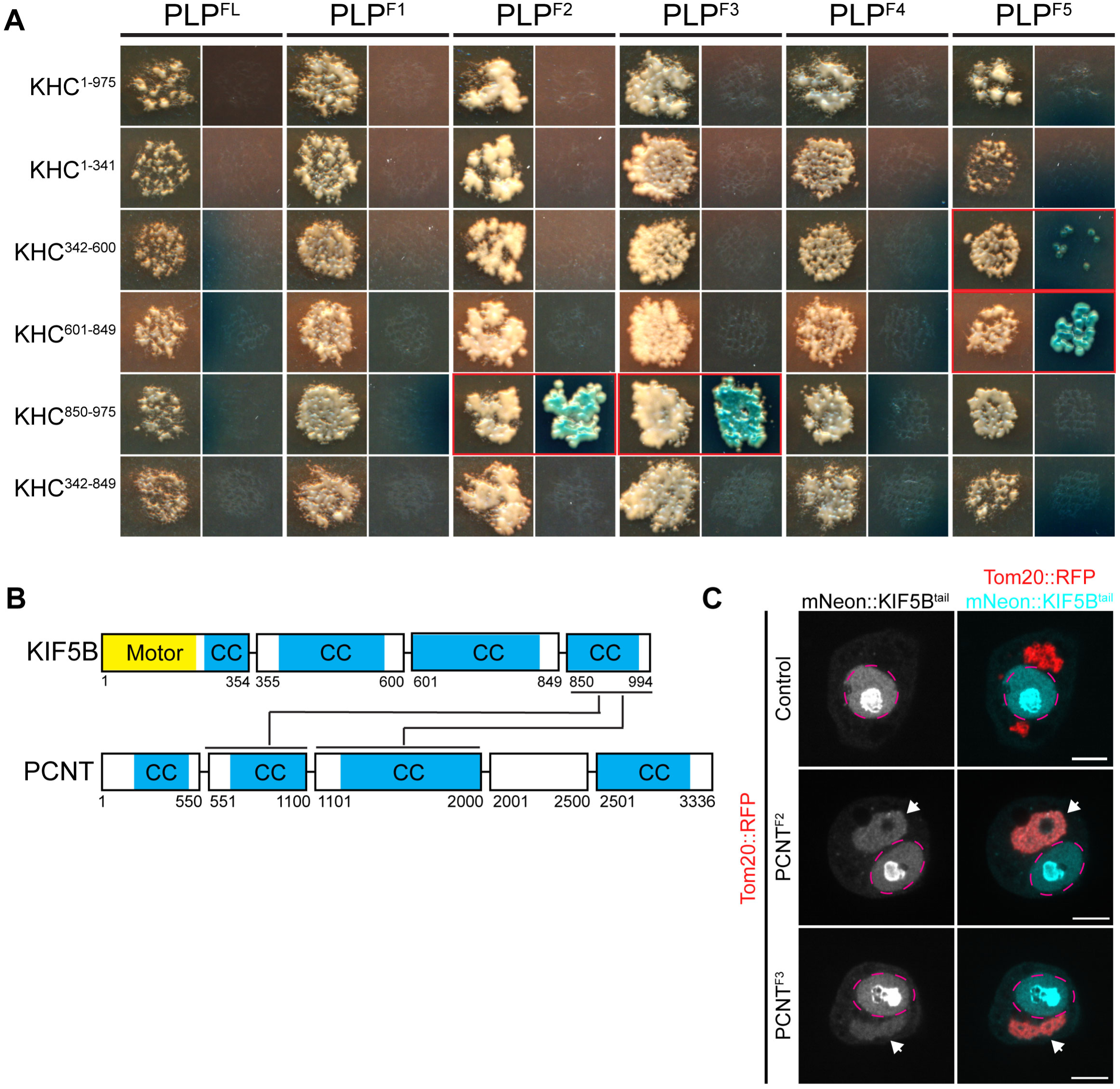
PLP-Kinesin-1 interaction is conserved. **A)** Images of mated yeast clones. For each PLP fragment the left column is DDO plates to select for both bait and prey. Right column is QDOXA plates to select for interaction. Red outlined clones indicate a positive interaction, identified by the growth of blue yeast colonies. **B)** Diagram illustrating interactions identified between human KIF5B and PCNT fragments, predicted coiled coils are highlighted in blue (CC).. **C)** Example images of S2 cells in which fragments of PCNT (Red) have been targeted to the mitochondria by a Tom20 mitochondrial targeting sequence. The KIF5B tail (aa850-994) was tagged with mNeonGreen. Note that KIF5B is only recruited to mitochondria in the presence of PCNT fragments (arrow heads) indicative of interaction. mNeon::KIF5B also accumulated in the nucleus (Magenta Dashed line). Scale bar: 5μm.

**Supplementary Figure 5:**
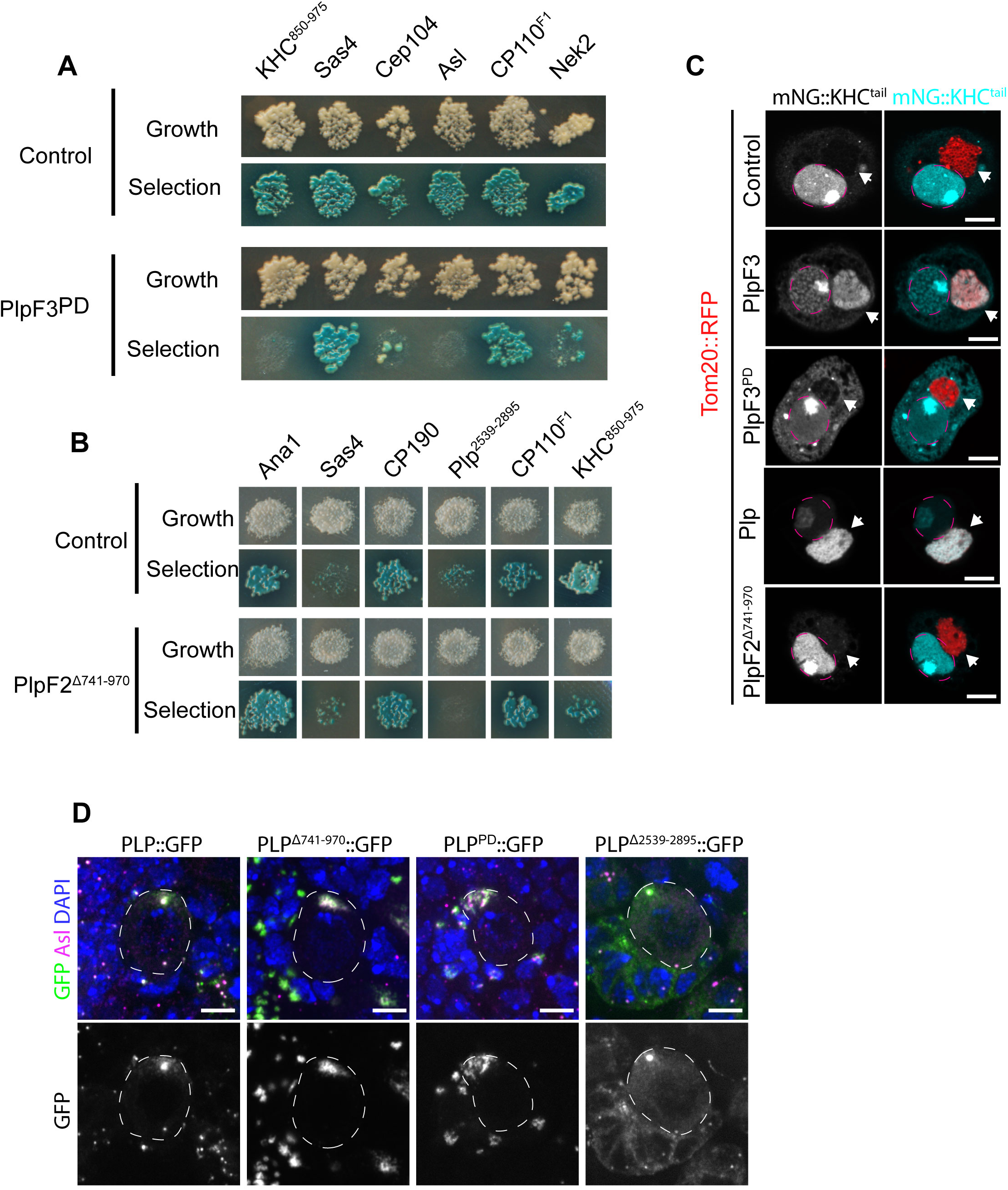
Disrupting the PLP-Kinesin-1 interaction. **A)** Yeast clones showing the PLP interactions disrupted by the PLP^PD^ mutation. Note that Asl interaction is also disrupted. Interaction was determined by the growth of blue clones on the selection plate (QDOXA). **B)** Yeast clones showing the interactions disrupted by deletion of PLP aa 741-970. The interaction with KHC^850-975^ appears weaker due to decreased growth. Note interaction with PLP^2539-2895^ is also disrupted. **C)** Validation of PLP-KHC interaction mutants by mitochondrial targeting assay, PLPF2 refers to PLP^584-1376^, PLPF3 refers to PLP^1377-1811^. PLP fragments were targeted to the mitochondria using the Tom20 mitochondrial localization sequence (red). The cargo binding tail of KHC (KHC^850-975^) was tagged with mNeon. Interaciton was determined by recruitment of mNeon::KHC^850-975^ to the mitochondria. (Arrow heads point to mitochondria, Magenta dashed line labels the nuclei). **D)** Over expression of PLP::GFP transgenes in NBs does not recapitulate endogenous localization. Left panel: the full length-rescue is no longer asymmetrically localized to the mother centriole and forms a large accumulation around the daughter centriole at the apical side of the cell. Center panels: The PLP-Kinesin-1 interaction mutants enhance the accumulation of protein to the apical side of the cell. Right panel: Deletion of PLP aa 2539-2895 does not prevent PLP centriolar localization. Scale bars: 5µm.

**Supplementary Figure 6:**
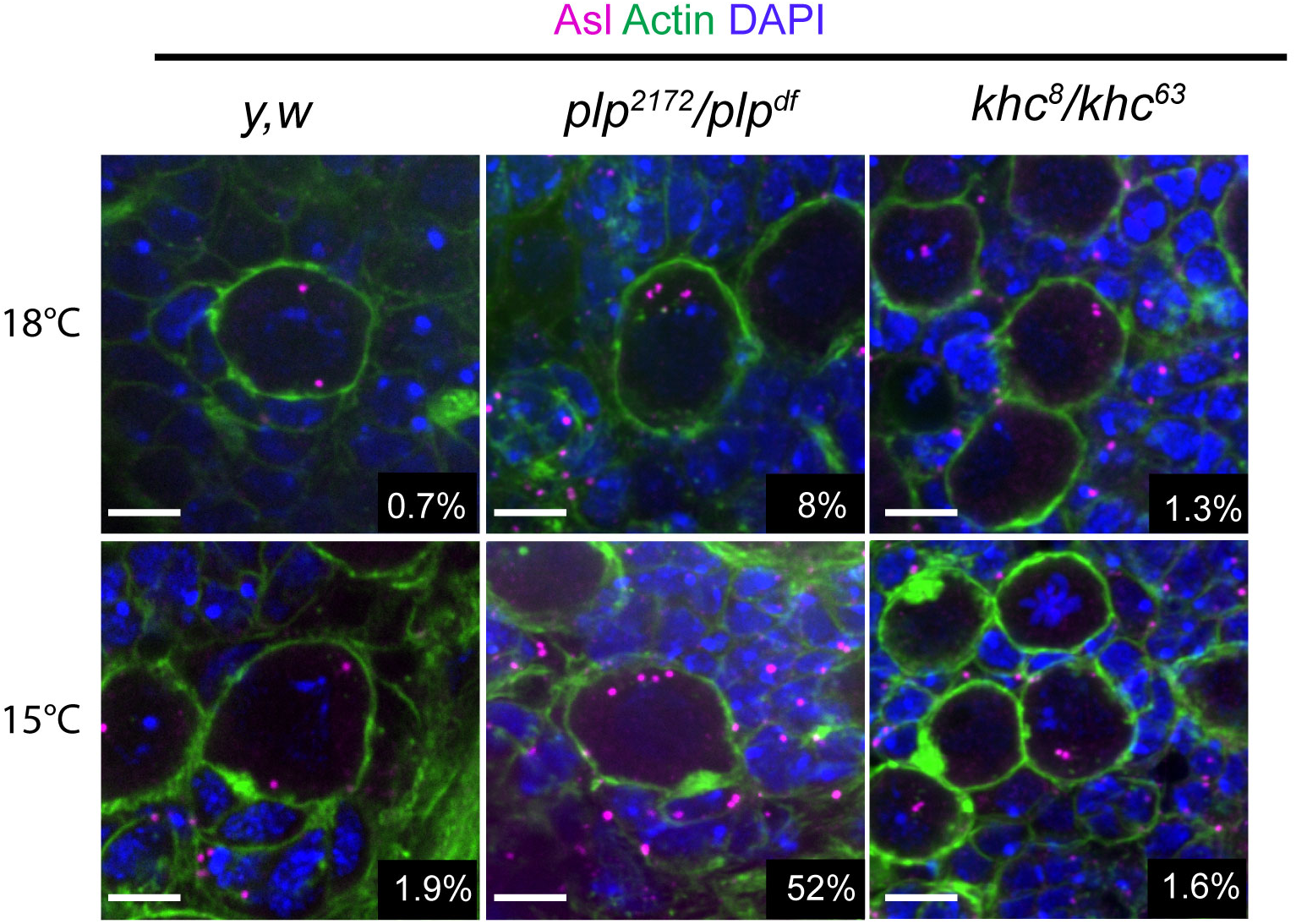
Increased supernumerary centrioles at low temperature is only observed in *plp* mutant NBs. Fixed NBs stained for Asterless (asl, magenta) to label centrioles, Phallodin (green) and DAPI (Blue). Centrioles were counted by counting the number of Asl positive puncta in each cell. Raising flies in lower temperatures only increased in the percentage of NBs carrying more than two Asl puncta in *plp* mutants (*plp^2172^/plp^df^).* 18°C: Control = 0.7%±0.8 (n=4 brains, 230 NBs), *plp* = 8.8%±3.6 (n=4 brains, 330 NBs), *khc* = 1.3%±2.7 (n=4 brains 235 NBs). 15°C: Control = 1.9%±1.5 (n=4 brains, 205 NBs), *plp* = 52%±6.7 (n=4 brains, 161 NBs), *khc* = 1.6%±2.3 (n=4 brains, 197 NBs). Data = Mean ± Standard Deviation. Scale bars: 5µm.

## Movies

**Movie 1: Centriole motility in *Drosophila* Neuroblasts.** Spinning disk confocal timelapse of an isolated neuroblast expressing Sas4::Neon (Cyan) and mCherry::Jupiter (Red). Note that one centriole migrates away from the apical side of the cell through interphase. Apical is up. Timestamp: hh:mm. Scalebar: 5µm

**Movie 2: Centrioles are motile in interphase S2 Cells.** Spinning disk confocal time-lapse of an Interphase *Drosophila* S2 cell expressing F-Tractin::RFP (Red) and GFP::PACT (centrioles, Cyan). Both centrioles are highly motile. Time stamp: mm:ss. Scale bar: 5µm.

**Movie 3: Centrioles are motile in interphase peripodial cells.** Spinning disk confocal timelapse of an interphase peripodial cell expressing UAS-Lifeact::RFP (Red) by AGIR-GAL4. Centrioles are labelled with ubi-GFP::Sas-6 (Cyan). Time stamp: mm:ss. Scale bar: 5µm.

**Movie 4: Centriole movement is independent of the Actin Cytoskeleton.** Spinning Disk confocal timelapse of peripodial cells expressing ubi-GFP::Sas6 to label centrioles. Wing discs were treated with DMSO (Left) or Latrunculin-A (Right). Note in both cases centrioles are highly motile. Time stamp: mm:ss. Scale bar: 5µm.

**Movie 5: Centriole movement is dependent upon the microtubule network but not microtubule dynamics.** Spinning disk confocal timelapse of interphase peripodial cells expressing ubi-Sas6::GFP to label centrioles. Top panels: Centrioles are highly motile after recovering in the presence of DMSO following exposure to ice (Left). Recovery from ice in the presence of Colcemid blocked centriole movement (Right). Bottom Panels: Treatment of wing discs with colchicine to inhibit microtubule dynamics does not block centriole movement (Right) compared to those treated with DMSO (Left). Time stamp: mm:ss. Scale bar: 5µm.

**Movie 6: Centrioles move on the microtubule network in peripodial cells.** Aireyscan microscopy timelapse showing centrioles (RFP::Pact, Cyan) moving along microtubules (GFP::Tubulin, Red). Timestamp: mm:ss. Scale bar: 2µm.

**Movie 7: Centrioles move on the microtubule network in neuroblasts.** Spinning disk confocal timelapse showing a mother centriole (Sas4::Neon, Cyan, Yellow arrowhead) moving away from the daughter centriole (Yellow asterisk) along the microtubule network (mCherry::Jupiter, Red) . Timestamp: mm:ss. Scale bar: 2µm.

**Movie 8: Centrioles switch between MTs at junctions.** Aireyscan timelapse of a peripodial cell labelled with SIR-Tubulin. The centriole (yellow arrowhead) moves along multiple microtubules, changing tracks at MT-MT junctions. Timestamp: mm:ss. Scale bar: 2µm.

**Movie 9: Centriole movement requires Kinesin-1.** Spinning disk confocal timelapse showing centriole (ubi-GFP::Sas6) movement following indicated knockdowns. RNAi was expressed under UAS promoter with the AGIR-GAL4. Time stamp: mm:ss. Scale bar: 5µm.

**Movie 10: Centriole movement is dependent upon PLP.** Spinning disk confocal timelapse showing centriole (ubi-GFP::Sas6) movement. Expressing UAS-PLPRNAi with Tub-GAL4 (Right) blocks centriole movement compared to controls (Left). Time stamp: mm:ss. Scale bar: 5µm.

**Movie 11: KHC and PLP^584-1811^ comigrate on microtubules *in vitro*. TIRF** timelapse showing comigration of PLP^584-1811^ (red) and KHC (green) on microtubules (Blue). Arrows indicate events of comigration. PLP molecules appear to precede KHC due to the slower multichannel imaging rate relative to the motility. Time: Seconds. Scalebar: 5µm.

**Movie 12: PLP Motility.** Movie showing movement of PLP^584-1811^ (red) on MTs in the presence of KHC. A maximum intensity projection of the KHC (blue) channel is used to highlight MT position. Time: seconds. Scalebar: 5µm.

**Movie 13: PLP-KHC interaction mutants fail to rescue centriole motility.** Spinning disk confocal timelapse of peripodial cells expressing indicated rescue constructs under UAS control by Engrailed-GAL4 in a *plp^-^* mutant background (*plp^2172^/Df(3L)Brd15*). Time stamp: mm:ss. Scale bar: 5µm.

**Movie 14: Centriole separation in neuroblasts**: Spinning disk confocal timelapses showing maximum projection of isolated neuroblasts expressing Sas4::Neon (Cyan) and mCherry::Jupiter (Red). Note centrioles (Cyan) do not migrate away from the apical side of the cell following PLP or KHC knockdown. Apical is up. Timestamp: hh:mm. Scalebar: 5µm

**Movie 15: Supernumerary centrioles following PLP knockdown is a result of incomplete prophase centrosome separation.** Spinning disk confocal timelapse showing maximum projection of an isolated neuroblast expressing UAS-PLPRNAi, Sas4::Neon (Centrioles, cyan) and mCherry::Jupiter (microtubules, red). As the cell enters mitosis the two centrosomes collapse to the apical spindle pole resulting in the segregation of both into the NB following mitosis. This then leads to four in the following cell cycle. Timestamp: hh:mm. Scalebar: 5µm.

**Movie 16: Polo::GFP asymmetry is disrupted following PLP or KHC knockdown.** Spinning disk confocal timelapse showing maximum projection of Polo::GFP in isolated neuroblasts. Movie pauses to show precocious activation of centrosomes following PLP (center) and KHC (right) knockdown relative to controls (left). Arrowheads indicate centrosome that gets segregated into the GMC. Timestamp: hh:mm. Scale bar: 5µm.

**Movie 17: Age dependent centrosome segregation is disrupted following PLP or KHC knockdown.** Spinning disk confocal timelapse showing maximum projection of isolated neuroblasts expressing ubi-Cbn::GFP (Daughter centriole, Cyan, Arrowhead) and UAS-mCherry::Jupiter (microtubules, red). Note that following PLP (center) or KHC (Right) knockdown the Cbn+ centriole ends up segregating to the basal side of the cell in mitosis rather than being retained by the NB at the apical side. Timestamp: hh:mm. Scale bar: 5µm.

## References

1. Agircan, F.G., E. Schiebel, and B.R. Mardin. 2014. Separate to operate: control of centrosome positioning and separation. Philosophical Transactions Royal Soc B Biological Sci. 369:20130461. doi:10.1098/rstb.2013.0461.

2. Ayloo, S., and E.L.F. Holzbaur. 2015. Chapter 4 Reconstitution of microtubule-based motility using cell extracts. Methods Cell Biol. 128:57–68. doi:10.1016/bs.mcb.2015.02.002.

3. Azimzadeh, J., and M. Bornens. 2007. Structure and duplication of the centrosome. J Cell Sci. 120:2139–2142. doi:10.1242/jcs.005231.

4. Bakhoum, S.F., and L.C. Cantley. 2018. The Multifaceted Role of Chromosomal Instability in Cancer and Its Microenvironment. Cell. 174:1347–1360. doi:10.1016/j.cell.2018.08.027.

5. Blasius, T.L., D. Cai, G.T. Jih, C.P. Toret, and K.J. Verhey. 2007. Two binding partners cooperate to activate the molecular motor Kinesin-1. J Cell Biology. 176:11–17. doi:10.1083/jcb.200605099.

6. Bolvar, J., J.R. Huynh, H. Lpez-Schier, C. Gonzlez, D.S. Johnston, and A. Gonzlez-Reyes. 2001. Centrosome migration into the Drosophila oocyte is independent of BicD and egl, and of the organisation of the microtubule cytoskeleton. Development. 128:1889– 1897. doi:10.1242/dev.128.10.1889.

7. Burakov, A., E. Nadezhdina, B. Slepchenko, and V. Rodionov. 2003. Centrosome positioning in interphase cells. J Cell Biology. 162:963–969. doi:10.1083/jcb.200305082.

8. Chen, C., and Y.M. Yamashita. 2021. Centrosome-centric view of asymmetric stem cell division. Open Biol. 11:200314. doi:10.1098/rsob.200314.

9. Ching, K., J.T. Wang, and T. Stearns. 2021. Long-range migration of centrioles to the apical surface of the olfactory epithelium. Biorxiv. 2021.10.12.464082. doi:10.1101/2021.10.12.464082.

10. Conduit, P.T., and J.W. Raff. 2010. Cnn Dynamics Drive Centrosome Size Asymmetry to Ensure Daughter Centriole Retention in Drosophila Neuroblasts. Curr Biol. 20:2187– 2192. doi:10.1016/j.cub.2010.11.055.

11. Crocker, J.C., and D.G. Grier. 1996. Methods of Digital Video Microscopy for Colloidal Studies. J Colloid Interf Sci. 179:298–310. doi:10.1006/jcis.1996.0217.

12. Dawe, H.R., H. Farr, and K. Gull. 2006. Centriole/basal body morphogenesis and migration during ciliogenesis in animal cells. J Cell Sci. 120:7–15. doi:10.1242/jcs.03305.

13. Djagaeva, I., D.J. Rose, A. Lim, C.E. Venter, K.M. Brendza, P. Moua, and W.M. Saxton. 2012. Three Routes to Suppression of the Neurodegenerative Phenotypes Caused by Kinesin Heavy Chain Mutations. Genetics. 192:173–183. doi:10.1534/genetics.112.140798.

14. Dujardin, D.L., and R.B. Vallee. 2002. Dynein at the cortex. Curr Opin Cell Biol. 14:44–49. doi:10.1016/s0955-0674(01)00292-7.

15. Friedman, D.S., and R.D. Vale. 1999. Single-molecule analysis of kinesin motility reveals regulation by the cargo-binding tail domain. Nat Cell Biol. 1:293–297. doi:10.1038/13008.

16. Fu, M., and E.L.F. Holzbaur. 2014. Integrated regulation of motor-driven organelle transport by scaffolding proteins. Trends Cell Biol. 24:564–574. doi:10.1016/j.tcb.2014.05.002.

17. Fuentealba, L.C., E. Eivers, D. Geissert, V. Taelman, and E.M.D. Robertis. 2008. Asymmetric mitosis: Unequal segregation of proteins destined for degradation. Proc National Acad Sci. 105:7732–7737. doi:10.1073/pnas.0803027105.

18. Gallaud, E., R. Caous, A. Pascal, F. Bazile, J.-P. Gagné, S. Huet, G.G. Poirier, D. Chrétien, L. Richard-Parpaillon, and R. Giet. 2014. Ensconsin/Map7 promotes microtubule growth and centrosome separation in Drosophila neural stem cells. J Cell Biology. 204:1111–21. doi:10.1083/jcb.201311094.

19. Gallaud, E., A.R. Nair, N. Horsley, A. Monnard, P. Singh, T.T. Pham, D.S. Garcia, A. Ferrand, and C. Cabernard. 2020. Dynamic centriolar localization of Polo and Centrobin in early mitosis primes centrosome asymmetry. Plos Biol. 18:e3000762. doi:10.1371/journal.pbio.3000762.

20. Galletta, B.J., C.J. Fagerstrom, T.A. Schoborg, T.A. McLamarrah, J.M. Ryniawec, D.W. Buster, K.C. Slep, G.C. Rogers, and N.M. Rusan. 2016. A centrosome interactome provides insight into organelle assembly and reveals a non-duplication role for Plk4. Nat Commun. 7:12476. doi:10.1038/ncomms12476.

21. Galletta, B.J., R.X. Guillen, C.J. Fagerstrom, C.W. Brownlee, D.A. Lerit, T.L. Megraw, G.C. Rogers, and N.M. Rusan. 2014. Drosophila pericentrin requires interaction with calmodulin for its function at centrosomes and neuronal basal bodies but not at sperm basal bodies. Mol Biol Cell. 25:2682–94. doi:10.1091/mbc.e13-10-0617.

22. Galletta, B.J., J.M. Ortega, S.L. Smith, C.J. Fagerstrom, J.M. Fear, S. Mahadevaraju, B. Oliver, and N.M. Rusan. 2020. Sperm Head-Tail Linkage Requires Restriction of Pericentriolar Material to the Proximal Centriole End. Dev Cell. 53:86–101.e7. doi:10.1016/j.devcel.2020.02.006.

23. Galletta, B.J., and N.M. Rusan. 2015. Methods in Cell Biology. Methods Cell Biol. 129:251– 277. doi:10.1016/bs.mcb.2015.03.012.

24. Ganem, N.J., S.A. Godinho, and D. Pellman. 2009. A Mechanism Linking Extra Centrosomes to Chromosomal Instability. Nature. 460:278–282. doi:10.1038/nature08136.

25. Gibson, M.C., D.A. Lehman, and G. Schubiger. 2002. Lumenal Transmission of Decapentaplegic in Drosophila Imaginal Discs. Dev Cell. 3:451–460. doi:10.1016/s1534-5807(02)00264-2.

26. Grieder, N.C., M. de Cuevas, and A.C. Spradling. 2000. The fusome organizes the microtubule network during oocyte differentiation in Drosophila. Development. 127:4253–4264. doi:10.1242/dev.127.19.4253.

27. Hammond, J.W., K. Griffin, G.T. Jih, J. Stuckey, and K.J. Verhey. 2008. Co-operative Versus Independent Transport of Different Cargoes by Kinesin-1. Traffic. 9:725–741. doi:10.1111/j.1600-0854.2008.00722.x.

28. Januschke, J., and C. Gonzalez. 2010. The interphase microtubule aster is a determinant of asymmetric division orientation in Drosophila neuroblasts. J Cell Biology. 188:693– 706. doi:10.1083/jcb.200905024.

29. Januschke, J., S. Llamazares, J. Reina, and C. Gonzalez. 2011. Drosophila neuroblasts retain the daughter centrosome. Nat Commun. 2:243. doi:10.1038/ncomms1245.

30. Januschke, J., J. Reina, S. Llamazares, T. Bertran, F. Rossi, J. Roig, and C. Gonzalez. 2013. Centrobin controls mother–daughter centriole asymmetry in Drosophila neuroblasts. Nat Cell Biol. 15:241–248. doi:10.1038/ncb2671.

31. Jord, A.A., N. Spassky, and A. Meunier. 2019. Motile ciliogenesis and the mitotic prism. Biol Cell. 111:199–212. doi:10.1111/boc.201800072.

32. Kapitein, L.C., E.J.G. Peterman, B.H. Kwok, J.H. Kim, T.M. Kapoor, and C.F. Schmidt. 2005. The bipolar mitotic kinesin Eg5 moves on both microtubules that it crosslinks. Nature. 435:114–118. doi:10.1038/nature03503.

33. Kelliher, M.T., Y. Yue, A. Ng, D. Kamiyama, B. Huang, K.J. Verhey, and J. Wildonger. 2018. Autoinhibition of kinesin-1 is essential to the dendrite-specific localization of Golgi outposts. J Cell Biology. 217:2531–2547. doi:10.1083/jcb.201708096.

34. Klebba, J.E., D.W. Buster, A.L. Nguyen, S. Swatkoski, M. Gucek, N.M. Rusan, and G.C. Rogers. 2013. Polo-like Kinase 4 Autodestructs by Generating Its Slimb-Binding Phosphodegron. Curr Biol. 23:2255–2261. doi:10.1016/j.cub.2013.09.019.

35. Krishnan, N., M. Swoger, M. Bates, J. Freshour, P.J. Fioramonti, A. Patteson, and H. Hehnly. 2021. Rab11 endosomes coordinate centrosome number and movement following mitotic exit. Biorxiv. 2021.08.11.455966. doi:10.1101/2021.08.11.455966.

36. Lerit, D.A., H.A. Jordan, J.S. Poulton, C.J. Fagerstrom, B.J. Galletta, M. Peifer, and N.M. Rusan. 2015. Interphase centrosome organization by the PLP-Cnn scaffold is required for centrosome function. J Cell Biol. 210:79–97. doi:10.1083/jcb.201503117.

37. Lerit, D.A., and N.M. Rusan. 2013. PLP inhibits the activity of interphase centrosomes to ensure their proper segregation in stem cells. J Cell Biology. 202:1013–22. doi:10.1083/jcb.201303141.

38. Liu, R., N. Billington, Y. Yang, C. Bond, A. Hong, V. Siththanandan, Y. Takagi, and J.R. Sellers. 2021. A binding protein regulates myosin-7a dimerization and actin bundle assembly. Nat Commun. 12:563. doi:10.1038/s41467-020-20864-z.

39. Lu, W., M. Winding, M. Lakonishok, J. Wildonger, and V.I. Gelfand. 2016. Microtubule-microtubule sliding by kinesin-1 is essential for normal cytoplasmic streaming in Drosophila oocytes. P Natl Acad Sci Usa. 113:E4995–5004. doi:10.1073/pnas.1522424113.

40. Martinez-Campos, M., R. Basto, J. Baker, M. Kernan, and J.W. Raff. 2004. The Drosophila pericentrin-like protein is essential for cilia/flagella function, but appears to be dispensable for mitosis. J Cell Biology. 165:673–683. doi:10.1083/jcb.200402130.

41. McClure, K.D., and G. Schubiger. 2005. Developmental analysis and squamous morphogenesis of the peripodial epithelium in Drosophila imaginal discs. Dev Camb Engl. 132:5033–42. doi:10.1242/dev.02092.

42. Métivier, M., B.Y. Monroy, E. Gallaud, R. Caous, A. Pascal, L. Richard-Parpaillon, A. Guichet, K.M. Ori-McKenney, and R. Giet. 2019. Dual control of Kinesin-1 recruitment to microtubules by Ensconsin in Drosophila neuroblasts and oocytes. Development. 146:dev171579. doi:10.1242/dev.171579.

43. Neighbors, B.W., R.C. Williams, and J.R. McIntosh. 1988. Localization of kinesin in cultured cells. J Cell Biology. 106:1193–1204. doi:10.1083/jcb.106.4.1193.

44. Pampalona, J., J. Januschke, P. Sampaio, and C. Gonzalez. 2015. Time-lapse recording of centrosomes and other organelles in Drosophila neuroblasts. Methods Cell Biol. 129:301–15. doi:10.1016/bs.mcb.2015.03.003.

45. Piel, M., P. Meyer, A. Khodjakov, C.L. Rieder, and M. Bornens. 2000. The Respective Contributions of the Mother and Daughter Centrioles to Centrosome Activity and Behavior in Vertebrate Cells. J Cell Biology. 149:317–330. doi:10.1083/jcb.149.2.317.

46. Piel, M., J. Nordberg, U. Euteneuer, and M. Bornens. 2001. Centrosome-Dependent Exit of Cytokinesis in Animal Cells. Science. 291:1550–1553. doi:10.1126/science.1057330.

47. Purohit, A., S.H. Tynan, R. Vallee, and S.J. Doxsey. 1999. Direct Interaction of Pericentrin with Cytoplasmic Dynein Light Intermediate Chain Contributes to Mitotic Spindle Organization. J Cell Biology. 147:481–492. doi:10.1083/jcb.147.3.481.

48. Ramat, A., M. Hannaford, and J. Januschke. 2017. Maintenance of Miranda Localization in Drosophila Neuroblasts Involves Interaction with the Cognate mRNA. Curr Biol. 27:2101–2111.e5. doi:10.1016/j.cub.2017.06.016.

49. Ramdas Nair, A., P. Singh, D. Salvador Garcia, D. Rodriguez-Crespo, B. Egger, and C. Cabernard. 2016. The Microcephaly-Associated Protein Wdr62/CG7337 Is Required to Maintain Centrosome Asymmetry in Drosophila Neuroblasts. Cell Reports. 14:1100– 1113. doi:10.1016/j.celrep.2015.12.097.

50. Rebollo, E., P. Sampaio, J. Januschke, S. Llamazares, H. Varmark, and C. González. 2007. Functionally Unequal Centrosomes Drive Spindle Orientation in Asymmetrically Dividing Drosophila Neural Stem Cells. Dev Cell. 12:467–474. doi:10.1016/j.devcel.2007.01.021.

51. Reiter, J.F., and M.R. Leroux. 2017. Genes and molecular pathways underpinning ciliopathies. Nat Rev Mol Cell Bio. 18:533–547. doi:10.1038/nrm.2017.60.

52. Rogers, G.C., N.M. Rusan, M. Peifer, and S.L. Rogers. 2008. A multicomponent assembly pathway contributes to the formation of acentrosomal microtubule arrays in interphase Drosophila cells. Mol Biol Cell. 19:3163–78. doi:10.1091/mbc.e07-10-1069.

53. Roque, H., S. Saurya, M.B. Pratt, E. Johnson, and J.W. Raff. 2018. Drosophila PLP assembles pericentriolar clouds that promote centriole stability, cohesion and MT nucleation. Plos Genet. 14:e1007198. doi:10.1371/journal.pgen.1007198.

54. Rusan, N.M., and M. Peifer. 2007. A role for a novel centrosome cycle in asymmetric cell division. J Cell Biology. 177:13–20. doi:10.1083/jcb.200612140.

55. Schoborg, T., A.L. Zajac, C.J. Fagerstrom, R.X. Guillen, and N.M. Rusan. 2015. An Asp-CaM complex is required for centrosome-pole cohesion and centrosome inheritance in neural stem cells. J Cell Biology. 211:987–98. doi:10.1083/jcb.201509054.

56. Sepulveda, G., M. Antkowiak, I. Brust-Mascher, K. Mahe, T. Ou, N.M. Castro, L.N. Christensen, L. Cheung, X. Jiang, D. Yoon, B. Huang, and L.-E. Jao. 2018. Co-translational protein targeting facilitates centrosomal recruitment of PCNT during centrosome maturation in vertebrates. Elife. 7:e34959. doi:10.7554/elife.34959.

57. Silkworth, W.T., I.K. Nardi, R. Paul, A. Mogilner, and D. Cimini. 2012. Timing of centrosome separation is important for accurate chromosome segregation. Mol Biol Cell. 23:401–411. doi:10.1091/mbc.e11-02-0095.

58. Singh, P., A. Ramdas Nair, and C. Cabernard. 2014. The Centriolar Protein Bld10/Cep135 Is Required to Establish Centrosome Asymmetry in Drosophila Neuroblasts. Curr Biol. 24:1548–1555. doi:10.1016/j.cub.2014.05.050.

59. Spassky, N., and A. Meunier. 2017. The development and functions of multiciliated epithelia. Nat Rev Mol Cell Bio. 18:423–436. doi:10.1038/nrm.2017.21.

60. Sun, F., C. Zhu, R. Dixit, and V. Cavalli. 2011. Sunday Driver/JIP3 binds kinesin heavy chain directly and enhances its motility. Embo J. 30:3416–3429. doi:10.1038/emboj.2011.229.

61. Swider, Z.T., R.K. Ng, R. Varadarajan, C.J. Fagerstrom, and N.M. Rusan. 2019. Fascetto interacting protein ensures proper cytokinesis and ploidy. Mol Biol Cell. 30:992–1007. doi:10.1091/mbc.e18-09-0573.

62. Tanenbaum, M.E., and R.H. Medema. 2010. Mechanisms of centrosome separation and bipolar spindle assembly. Dev Cell. 19:797–806. doi:10.1016/j.devcel.2010.11.011.

63. Tang, N., and W.F. Marshall. 2012. Centrosome positioning in vertebrate development. J Cell Sci. 125:4951–4961. doi:10.1242/jcs.038083.

64. Tinevez, J.-Y., N. Perry, J. Schindelin, G.M. Hoopes, G.D. Reynolds, E. Laplantine, S.Y. Bednarek, S.L. Shorte, and K.W. Eliceiri. 2017. TrackMate: An open and extensible platform for single-particle tracking. Methods. 115:80–90. doi:10.1016/j.ymeth.2016.09.016.

65. Tozer, S., C. Baek, E. Fischer, R. Goiame, and X. Morin. 2017. Differential Routing of Mindbomb1 via Centriolar Satellites Regulates Asymmetric Divisions of Neural Progenitors. Neuron. 93:542–551.e4. doi:10.1016/j.neuron.2016.12.042.

66. Tripathi, A., C. Bond, J.R. Sellers, N. Billington, and Y. Takagi. 2021. Myosin-Specific Adaptations of In vitro Fluorescence Microscopy-Based Motility Assays. J Vis Exp. doi:10.3791/62180.

67. Varadarajan, R., and N.M. Rusan. 2018. Bridging centrioles and PCM in proper space and time. Essays Biochem. 62:793–801. doi:10.1042/ebc20180036.

68. Wang, X., N. Le, A. Denoth-Lippuner, Y. Barral, and R. Kroschewski. 2016. Asymmetric partitioning of transfected DNA during mammalian cell division. Proc National Acad Sci. 113:7177–7182. doi:10.1073/pnas.1606091113.

69. Winding, M., M.T. Kelliher, W. Lu, J. Wildonger, and V.I. Gelfand. 2016. Role of kinesin-1– based microtubule sliding in Drosophila nervous system development. Proc National Acad Sci. 113:E4985–E4994. doi:10.1073/pnas.1522416113.

70. Young, A., J.B. Dictenberg, A. Purohit, R. Tuft, and S.J. Doxsey. 2000. Cytoplasmic Dynein-mediated Assembly of Pericentrin and γ Tubulin onto Centrosomes. Mol Biol Cell. 11:2047–2056. doi:10.1091/mbc.11.6.2047.

